# Enhancer-driven local 3D chromatin domain folding modulates transcription in human mammary tumor cells

**DOI:** 10.1101/2023.10.17.562690

**Authors:** Silvia Kocanova, Flavien Raynal, Isabelle Goiffon, Betul Akgol Oksuz, Davide Baú, Alain Kamgoué, Sylvain Cantaloube, Ye Zhan, Bryan Lajoie, Marc A. Marti-Renom, Job Dekker, Kerstin Bystricky

**Affiliations:** Molecular, Cellular and Developmental biology unit (MCD); Centre de Biologie Integrative (CBI); University of Toulouse; UPS; CNRS; 31062 Toulouse, France; Institut Universitaire de France (IUF); Department of Systems Biology; University of Massachusetts Chan Medical School; Worcester MA, 01605, USA; Howard Hughes Medical Institute, Chevy Chase, MD, USA; Centre Nacional d’Anàlisi Genòmica (CNAG), Barcelona, Spain; Genome Biology Program, Centre de Regulació Genòmica (CRG), Barcelona, Spain; Universitat Pompeu Fabra (UPF), 08002 Barcelona, Spain; Institució Catalana de Recerca i Estudis Avançats (ICREA), Barcelona, Spain

## Abstract

The genome is organized in functional compartments and structural domains at the sub-megabase scale. How within these domains interactions between numerous cis-acting enhancers and promoters regulate transcription remains an open question. Here, we determined chromatin folding and composition over several hundred kb around estrogen responsive genes in human breast cancer cell lines following hormone stimulation. Modeling of 5C data at 1.8 kb resolution was combined with quantitative 3D analysis of multicolor FISH measurements at 100 nm resolution and integrated with ChIP-seq data on transcription factor binding and histone modifications. We found that rapid estradiol induction of the progesterone gene (PGR) expression occurs in the context of pre-existing, cell type specific chromosomal architectures encompassing the 90 kb PGR coding region and an enhancer-spiked 5’ 300 kb upstream genomic region. In response to estradiol, interactions between estrogen-receptor α (ERα) bound regulatory elements are re-enforced. While initial enhancer – gene contacts coincide with RNA Pol 2 binding and transcription initiation, sustained hormone stimulation promotes ER accumulation creating a regulatory hub stimulating transcript synthesis. In addition to implications for estrogen receptor signaling, we uncover that preestablished chromatin architectures efficiently regulate gene expression upon stimulation without the need for de-novo extensive rewiring of long-range chromatin interactions.

## Introduction

Rapid cellular responses to external stimuli rely on regulation of gene expression. Among numerous steps required for this response, the spatial organization of the genome is known to modulate DNA accessibility to the transcriptional machinery and to promote contacts between genes and distant regulatory DNA elements such as enhancers (Gheldof et al., 2010; Kooren et al., 2007; Oudelaar et al., 2021; Paliou et al., 2019; Smith et al., 2016). In the past two decades, different levels of three-dimensional (3D) folding of the genome have been described thanks to ever improving technologies from population based contact frequencies to single cell imaging at high resolution (Gibcus and Dekker, 2013; Kim and Shendure, 2019; van Steensel and Furlong, 2019). Entire chromosomes adopt differential conformations as a function of transcriptional competence, for example between the active and inactive X-chromosomes (Boninsegna et al., 2022). All chromosomes are organized into active and inactive compartments composed of mega-base domains, and, at the scale of 10-100s of kb, can form topologically associating domains (TADs). TADs reflect areas of increased contact probabilities between DNA elements, but their intrinsic organization is highly complex, variable between cells in the population, and their relevance subject to debate (Sikorska and Sexton, 2020). There is no doubt, however, that numerous genes and their regulatory elements are located within a single TAD, and that TAD boundaries reduce the probability of long-range contact between elements and genes located on different sides. How this relates to controlled transcriptional activity remains to be fully understood.

3D chromatin folding reorganizes during differentiation and development reflecting changes in transcriptional activity (Tan et al., 2021). Estradiol (E2) signaling is a well-studied paradigm for transcriptional regulation in mammalian cells. This hormone exerts essential pleiotropic actions during development and differentiation and is best known for its role in the function of reproductive tissues of both males and females. A major role of estrogens is to modulate the transcriptional status of target genes, with these actions being transduced through a specific nuclear receptor, the Estrogen Receptor (ERα). ERα is a major determinant of tumor growth in about 80% of breast cancers in which it controls cell cycle genes such as Cyclin D1 (CCND1) as well as differentiation genes such as the progesterone receptor gene (PGR) and the estrogen receptor α coding gene itself (ESR1). Estrogens and hormones in general, control transcription of hundreds of genes in the eukaryotic nucleus. It was shown that changes in gene expression are accompanied by modifications in global genome structure in ER-expressing cells (Le Dily et al., 2014; Zhang et al., 2020). Yet, individual TADs were maintained during hormone-induced activation. Le Dily *et al*. proposed that the formation of structural regulons could promote coordinated expression of several genes. Not all of these genes would directly be estrogen regulated since ERα target genes do not need to colocalize to be activated (Kocanova et al., 2010a). Numerous estrogen receptor binding sites (ERBSs) are not only present at some promoters of ERα target genes but are also found located at 10-100 kb distance from target genes. A role in attracting cofactors to connect these distant elements and target genes has been proposed for a subset of ER-regulated genes (Kininis et al., 2007) and Chia-PET analysis suggests increased contacts (Fullwood et al., 2009). We thus asked if TADs are gene specific, reflecting and/or contributing to regulation of the gene and other regulatory elements within a domain.

Here, we integrate data from high resolution 5C, 3D FISH, ChIP-seq and computational modeling to analyze structural features of genomic domains containing several ERα target genes in two human breast cancer cell lines. Integration of data from such orthogonal experimental approaches enable establishing models of nuclear organization and defining statistically relevant structure/function relationships (Abbas et al., 2019; Boninsegna et al., 2022; Nir et al., 2018; Szabo et al., 2020). We show that folding of ERα target gene domains differs between the silent and the activatable form of the ESR1 and PGR genes. In ERα positive MCF7 cells, the pre-established domain conformation reorganizes upon E2 induced stimulation bringing distant ERα bound enhancer elements in proximity of the gene body.

## Results

### High resolution 3D maps of genomic domains encompassing estrogen sensitive genes are cell type specific

To determine whether the 3D organization of chromatin domains correlates with transcriptional status, we generated 5C chromatin interaction maps of ERα target genes whose transcriptional status differs in two human breast cancer cell lines with distinct tumor origins, MCF7 and MDA-MB-231 (Fig. 1A). These cell lines are representative of breast cancer (BC) types: MCF7 cells express the estrogen receptor alpha (ERα+) and their growth is hormone dependent. MDA-MB-231 cells are triple negative, hence do neither express the estrogen receptor alpha (ERα-), the progesterone receptor (PGR), nor the human epidermal growth factor receptor 2 (Her2) and their growth is independent of hormone. We selected 0.6 to 1.3 Mb domains around four ER-regulated genes: the estrogen receptor gene (ESR1) located on chromosome 6, the growth regulation by estrogen in breast cancer 1 gene (GREB1) located on chromosome 2, the cyclin D1 gene (CCND1) and the progesterone receptor gene (PGR) on chromosome 11, and determined their 3D conformation using 5C (Dostie et al., 2006) in MCF7 and MDA-MB-231 cell lines (Fig. 1B). Genes were chosen based on the transcriptional status and estrogen-responsiveness (Giamarchi et al., 1999; Honkela et al., 2015). Expression levels of the analyzed genes were confirmed by RT-qPCR (Fig. S1A). PGR and ESR1 were inducible in MCF7 cells and silent in MDA-MB-231 cells. GREB1 and CCND1 were constitutively transcribed in MDA-MB-231 cells, and were hormone-inducible in MCF7 cells (Fig. S1A).

**Figure 1:**
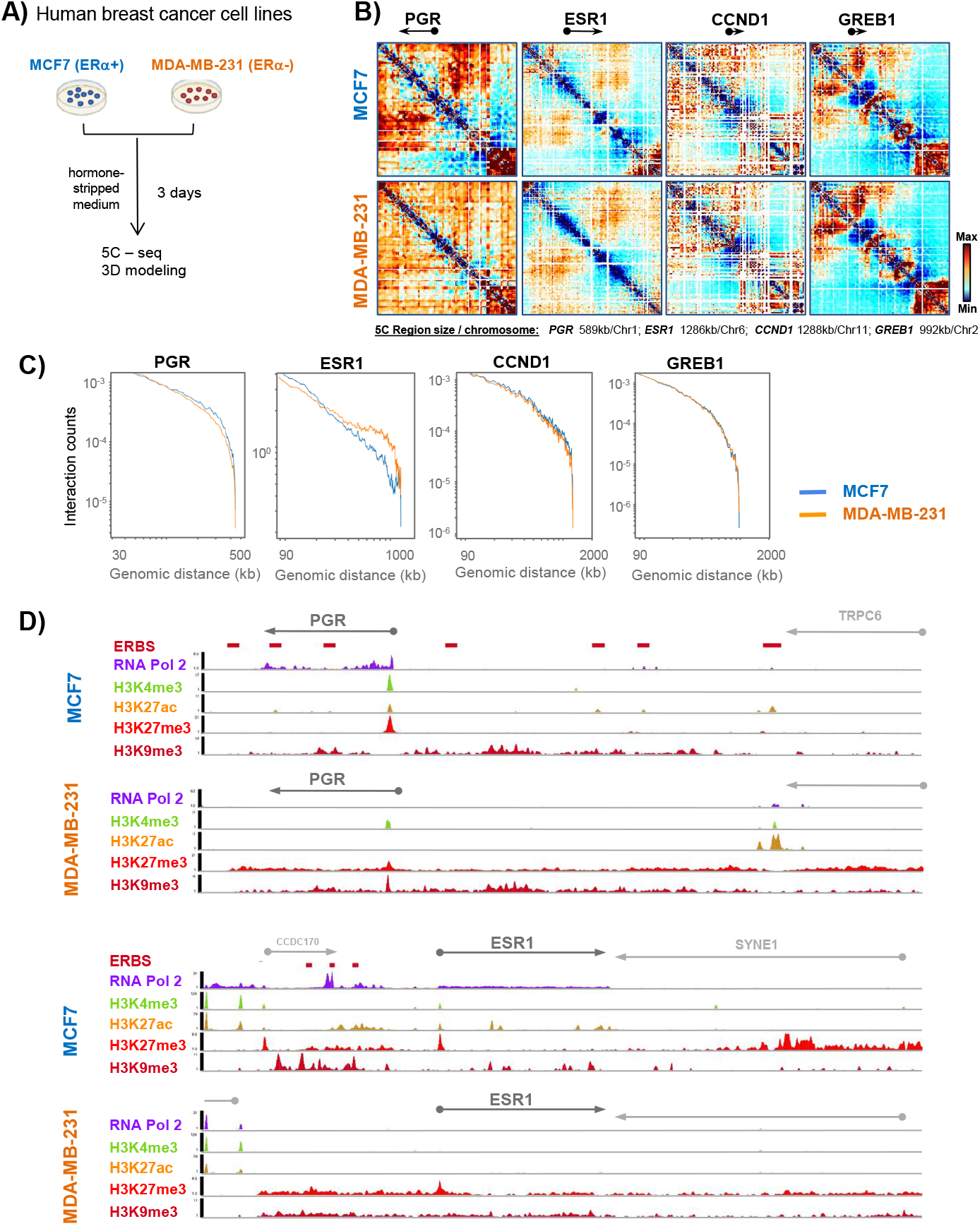
Pre-established genome organization of estrogen regulated gene domains reflects transcriptional status. A) Human breast cancer cell preparations and experimental flow used to interrogate 3D genome organization in cell lines expressing or not ER . B) Interaction frequency 5C heat maps at 1.8 kb resolution of 0.2 to 1.3 Mbp domains surrounding the PGR, ESR1, CCND1 and GREB1 genes in MCF7 and MDA-MB-231 cells. C) Genomic interaction counts of the selected gene domains decayed by genomic distance, comparison between MCF7 (blue line) and MDA-MB-231 (red line) cells. D) Chromatin landscape of the PGR and the ESR1 gene domains in MCF7 and MDA-MB-231 cells, from ENCODE (ENCODE Project Consortium, 2012; Guertin et al., 2014; Luo et al., 2020).

We used Chromosome Conformation Capture Carbon Copy (5C) with an alternating primer design (Dostie et al., 2006; Kim and Dekker, 2018) to assess contact frequencies at a resolution of 1.8 kb from cells growing in hormone-stripped medium for 3 days (Fig. 1A). 5C contact frequency maps revealed a high degree of similarity of the overall organization of the studied domains between the two phenotypically distinct cell lines. Within the CCND1 and GREB1 gene domains, only a few architectural features were cell type specific in agreement with the fact that these genes are expressed in both cell lines (Fig. 1B, Fig. S1). In contrast, significant conformational differences were detected for the PGR and ESR1 gene domains. PGR and ESR1 were silent in MDA-MB-231 cells, but were transcriptionally active in MCF7 cells (Fig. S1). In MCF7 cells, the PGR domain features two TADs with a clear boundary at the 3’end of the TRPC6 gene. TRPC6 codes for a transient receptor channel complex overexpressed in BC cells as compared to non-tumorous cell lines (Jardin et al., 2018) (Fig.S1B). This TAD organization was not present in MDA-MB-231 cells in which numerous weak long contact frequencies characterize the entire domain and no TAD boundary was detected (Fig. S1B). The PGR gene located within a 450 kb region which corresponds to the first TAD (TAD1), revealed slightly stronger interaction frequencies at long distances in MCF7 compared to MDA-MB-231 cells (Fig. 1C, S1B). The ESR1 gene domain featured greater long-distance interactions in MDA-MB-231 cells compared to MCF7 cells (Fig. 1C, S1C green arrowhead), interactions which appeared to stem from loci flanking the gene itself and the 5’ end of SYNE1 (Fig. 1B). A domain boundary is present within the SYNE1 gene. SYNE1 is rarely transcribed in MCF7 cells (proteinatlas.org). In MDA-MB-231 cells, SYNE1 is expressed and it is possible that transcription may lead to loss of this boundary, allowing increased interactions between the 5’ part of SYNE1 with the rest of the locus including ESR1. 5C contact maps and interaction counts for the control gene PUM1 (Kılıç et al., 2014) were similar in MCF7 and MDA-MB-231 cells (Fig. S1D and S1E).

To further explore the chromatin landscape of the four ER-regulated genes, we analyzed ChIP-seq data (Guertin et al., 2014) for histone post-translational modifications (PMT) and RNA polymerase 2 (RNA Pol2) for the two cell lines. The data show that RNA Pol2 was present at all representative genes in MCF7 but absent in MDA-MB-231 cells, except at the CCND1 gene which was constitutively active in MDA-MB-231 (Fig. 1D and S1A, S1F, and S1G). The H3K27ac and H3K4me3 chromatin modifications were present at the TSS and/or ERBS of ER-dependent genes in MCF7 but largely absent at genes silent in MDA-MB-231 (Fig. 1D and S1A, S1F, and S1G). The H3K27me3 and H3K9me3 repressive chromatin marks largely covered the gene bodies and surrounding regulatory domains in MDA-MB-231 (Fig. 1D, S1F, and S1G). Seven estrogen receptor binding sites (ERBS) exist within the TAD1 - PGR domain including the 100 kb coding region and an ~300 kb upstream regulatory, intergenic region (Fig. 1D). ERBS are reminiscent of enhancers and their chromatin was modified by H3K27ac in MCF7, a PMT absent at the PGR ERBSs in MDA-MB-231, except at the last, the 7^th^, ERBS (Fig. 1D). Contacts seen in 5C appeared to occur mainly between the PGR gene body and distal up-stream enhancers but not between enhancers and the transcription start site (TSS) of the gene itself (Fig. S1B). The ESR1 gene domain comprises three ER binding sites (ERBS) upstream of the ESR1 gene. ERBS contacted each other in MCF7 cells but these interactions were not detected in the hormone-independent MDA-MB-231 cells (Fig. S1C). H3K27me3 peaks were found at all promoters in both cell lines while this mark only covered the gene bodies of ESR1 and PGR silenced in MDA-MB-231cells (Fig. 1D), consistent with broader contact frequencies (Fig. 1B). The cell type specific contact maps suggested that the 3D conformation of these gene domains relates to transcriptional competence.

### Pre-established 3D domain architecture stabilizes in response to estradiol stimulation of PGR transcription

To further elucidate how chromatin folding is linked to transcriptional status, we focused on the PGR gene. We generated 5C contact maps of the PGR domain from MCF7 cells grown in hormone-stripped medium prior to and after 45 minutes and 3 hours of adding 100 nM E2 (Fig. 2A and 2B). Transcription of PGR increased 3 fold after 3 hour of incubation in the presence of E2 (Dalvai and Bystricky, 2010; Kocanova et al., 2010a). We found that the overall domain architecture, in particular the boundary next to the 3’end of the TPCR6 gene, was maintained following hormone addition (Fig. 2B). Visual inspection of 5C contact maps suggested that chromatin interactions between the TSS of the PGR and the gene body as well as its down-stream region (ERBS1, ERBS2 and ERBS3) were lost 3h after E2 stimulation (Fig. 2B, red arrowheads). In addition, upon early response to E2 several distinct contacts were reinforced between the TSS and proximal up-stream region of the PGR notably between TSS-ERBS4 (Fig. 2B and 2D). Gain of contacts between distal up-stream enhancers (ERBS5, ERBS6 and ERBS7) were also observed upon 3 hours of E2 stimulation (Fig. 2B, green arrowhead).

**Figure 2:**
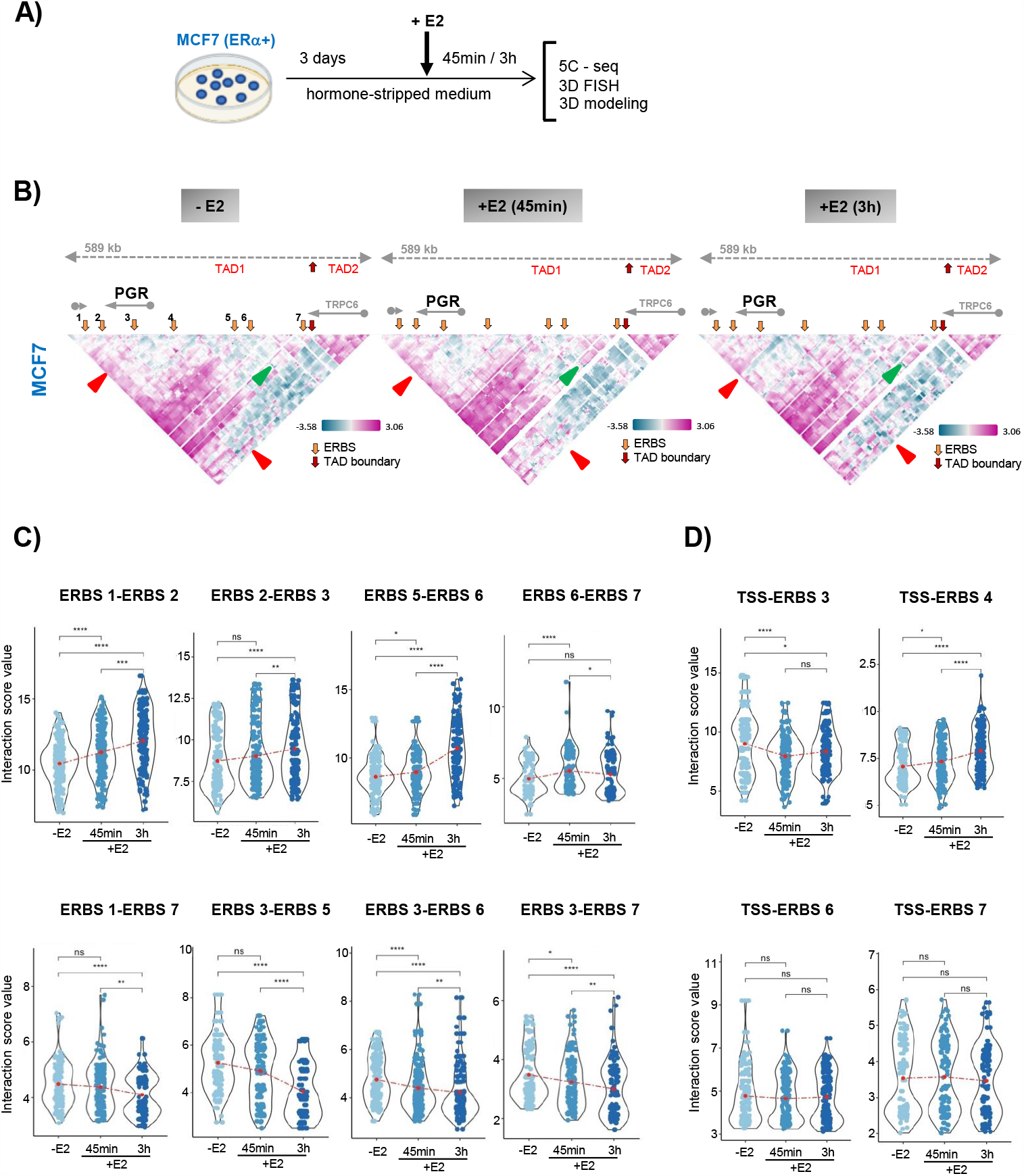
Ground-state domain architecture of the PGR domain stabilizes, and contact frequencies between regulatory elements are modulated, in response to estradiol signaling. A) Experimental flow used to interrogate 3D genome organization in MCF7 cells treated or not with E2. B) Interaction frequency 5C heat maps of the PGR gene domain in estrogen starved (-E2) and stimulated (+ E2) for 45 min and 3 h MCF7 cells. TAD boundaries do not change in MCF7 cells (red arrows) and ESRBs are indicated (yellow arrows). Increased (green arrowhead) and lost (red arrowhead) contact frequencies are indicated. C) Quantification of ERBS-ERBS interactions is calculated from 5C heat maps. Representing the gain (first row) and lost (second row) of interactions between specific ERBSs. D) Quantification of ERBS-TSS interactions showing variation in interaction with different ERBSs.

To validate the specific interactions, we quantified contact frequencies between all ERBSs and between TSS-ERBSs (Fig. 2C and 2D). We found that the interactions between ERBS3-ERBS5, ERBS3-ERBS6 and ERBS3-ERBS7 were slightly reduced at 45 min before declining at 3h. A drop in interactions was also detected at the boundaries of the TAD1 domain, between ERBS1-ERBS7 (Fig. 2C, lower panel). Two downstream (ERBS1-ERBS2 and ERBS2-ERBS3) and the upstream (ERBS5-ERBS6) regions established interactions at 45 min which were reinforced at 3 h. Moreover, interactions between distal upstream enhancers of the PGR gene (ERBS6-ERBS7) appeared as soon as 45 min of E2 stimulation and remained stable at 3h E2 (Fig. 2C, upper panel). Notably, the promoter (TSS) of the gene did not form detectable contacts with ERBSs except with ERBS4 (Fig. 2D). We noticed that TSS-ERBS4 contacts were strongly reinforced after 45 min E2 and, similar to ERBS1-ERBS2 and ERBS5-ERBS6, this interaction persisted over 3h E2 (Fig. 2C and 2D). Immediately after 45min of E2 addition contacts between TSS-ERBS3, the regulatory element located within the gene body, were lost (Fig. 2D). We conclude that enhancer interactions within the regulatory regions of PGR are modulated in an early response to hormone addition and reinforced during prolonged stimulation.

### Estrogen activation increases distal enhancer-enhancer interactions, and stabilizes folding of the PGR domain

To investigate the chromatin architecture of the PGR domain *in situ*, we analyzed the spatial relation between the PGR promoter and its enhancers (ERBSs) by 3D DNA FISH (Kocanova et al., 2018). Fosmid probes were selected according to availability corresponding to ERBSs and the TSS of the PGR gene (Fig. 3A). We performed 3D DNA FISH on MCF7 cells prior to and after 45 min and 3 h of 100 nM E2 stimulation and we measured the inter-probe distances using homemade script (plug in) running on ImageJ (Fig. 3B). 3D distance measurements between pairs of fosmid probe signals were plotted (Fig. 3C and S2A). In general, upon E2 activation, we noted large variations in distances measured. Distances spread from 20 nm to 1000 nm for fosmids separated by genomic distances from 78kb to 227kb (Fig. 3C and S2A). Variations of inter-probe distances -/+ addition of E2 were not significant population-wide. When focusing on distances ≤ 200 nm, which represent instances detectable by 5C (Giorgetti and Heard, 2016), inter-probe measures between Fos4-Fos5 and Fos3-Fos4, located in the upstream domain, 100kb and 250kb from the PGR-TSS increased significantly (Fig. 3D and S2B). For example, for Fos4-Fos5 and Fos3-Fos4, the proportion of measurements ranged from 4% and 19% in E2-non-treated cells to 23% and 29% in 3h E2 treated cells, respectively (Fig. 3D and S2B). In contrast, distances ≤ 200 nm significantly diminished between Fos3, covering the TSS, and Fos2, a fosmid probe located within the PGR gene body. We observed a decrease of short inter-probe distances between Fos2-Fos3 from 24% in untreated cells compared to 14% after 3h E2 stimulation (Fig. 3D). Distances between Fos3 (TSS) and Fos1 located in the downstream domain of the PGR gene were also significantly changed, with 50% of the contacts at distances ≤ 200 nm lost 3h after E2 stimulation (Fig. 3D). Inter-probe distances <200 nm at the PUM1 control locus did not vary between E2 untreated and treated MCF7 cells (Fig. S2C and S2D). Variations in inter-probe distances from 3D DNA FISH observations correlate with changes in contact frequencies seen in the same areas in 5C matrices. Both by imaging and 5C, we measured significant changes of interaction between the TSS and the upstream and downstream regulatory regions of PGR (Fig. 2B, 2C and 2D; Fig. 3; Fig. S2A, and S2B).

**Figure 3:**
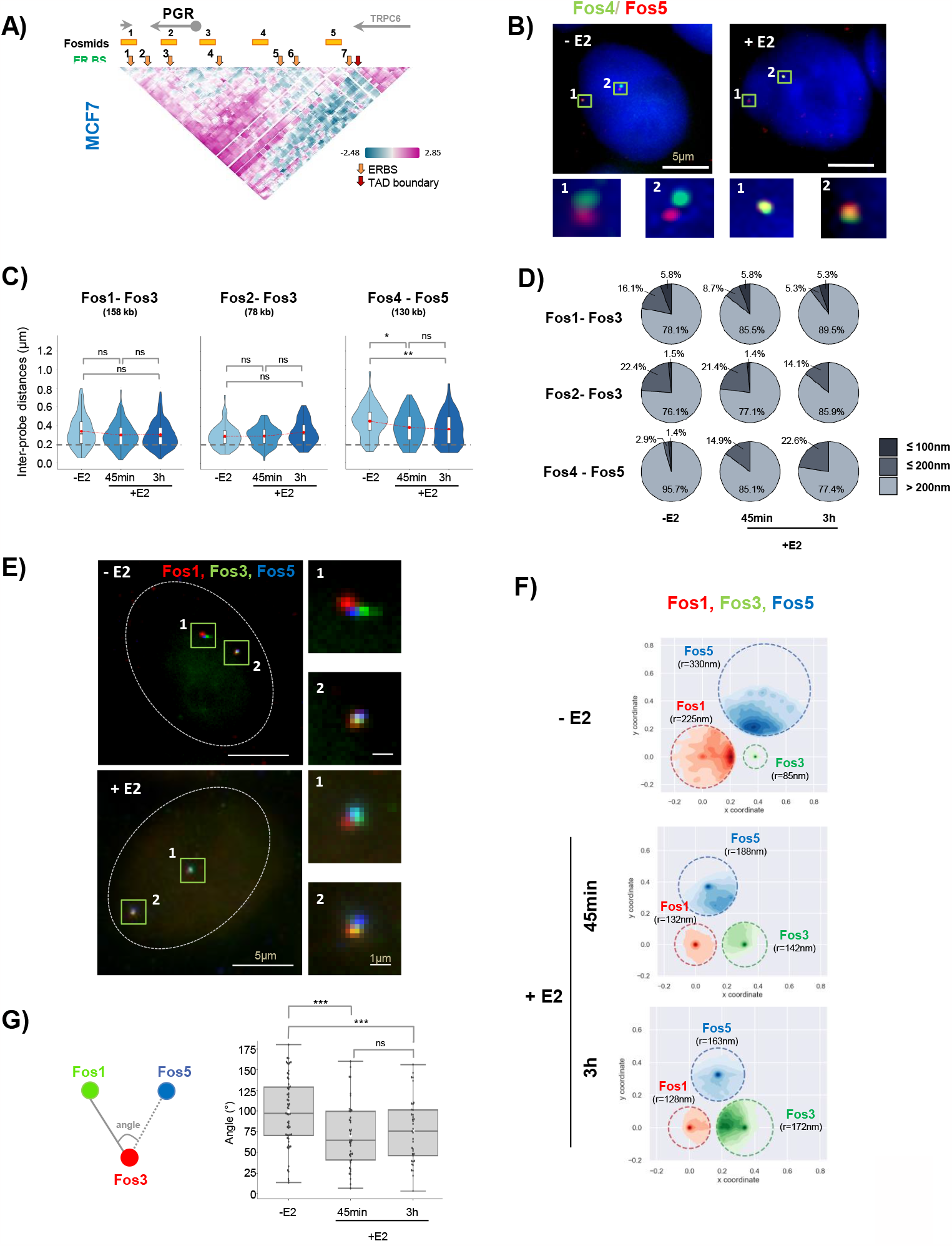
Interactions between regulatory elements of the PGR gene domain are re-enforced in response to estradiol signaling in MCF7 cells. A) Genomic position of fosmid probes used for 3D DNA FISH analysis. B) Representative images from 3D imaging of dual-fosmid labeled and DAPI co-stained nuclei of MCF7 cells stimulated (+E2) or not (-E2) with 100 nM estradiol. Fos4 labeled with Alexa488 (green), Fos5 in red (labeled with Atto 647). Maximal projection of 3 planes is presented (0.2 µm per single plane). See MM for details. Scale bar 5µm. C) The violin plots representing inter-probe distances for different pairs of fosmids (n= 80-130 nuclei). Fisher’s test: *P-values: >0.05 (ns), <0.05 (*), <0.01(**), <0.001 (***), <0.0001 (****). D) The pie charts illustrating the proportion of inter-probe distances > 200 nm, interval from 200-100nm (≤ 200 nm) and interval from 0 -100 nm (≤100 nm) for different pairs of fosmids in MCF7 cell stimulated or not by E2. Fisher’s test was used for statistics. E) Representative images from 3D imaging of triple-fosmid-labeled and DAPI co-stained nuclei of MCF7 cells stimulated (+E2) or not (-E2) with 100 nM estradiol. Maximal projection of 3 planes for Fos1 (in red), Fos3 (in green) and Fos5 (in blue) are presenting. Scale bar 5µm. F) Single cell analysis of relative position of three loci simultaneously representing as survival zone distribution of Fos1, Fos3 and Fos5. See MM for details. G) Angle around the Fos3 measured from mutual position of the three loci in single cell. Fisher’s test was used for statistics.

The conformational changes within the PGR domain indicated that the upstream regulatory region folds back upon itself and the PGR gene body upon transcription induction of PGR by E2. To characterize folding behavior of multiple connected mobile genomic loci we developed an analysis, called 3loci, based on triangulation of relative distances between three labeled sites. 3loci is based on measurements of the distribution of 3D distances between three loci computed in single cells. Within the plane defined by the three loci, we consider that each locus moves in a circular region with radius *R*. So, ***Sz****=****Sz****(R)* defining a 2D survival zone (Sz) with one degree of freedom. Using the measured distances, inverse mathematical modeling predicts each locus’ distribution inside survival zones (materials and methods). Survival zone radii of loci correlate with relative freedom of movement of one locus with respect to the two others (Lassadi et al., 2015). Here, we used three fosmids (Fig. 3E) hybridizing to the PGR gene and two ERBS within the up- and downstream regulatory regions simultaneously. In each nucleus, three distances were measured in 3D and aggregated to determine their survival zones. In cells treated with E2, survival zones of the two labeled loci (Fos1 and Fos5) surrounding the PGR gene were largely reduced compared to the zones these segments can explore in hormone deprived cells. In contrast, the centrally located Fos3 locus which corresponds to the TSS region of the PGR seemed to become more dynamic after transcription activation by E2. Upon activation, the distal upstream region (Fos5) folds toward the PGR gene body. Its freedom of movement is reduced two-fold (r=330 nm to 163 nm, Fig. 3F). The dynamics were decreased and this decrease was maintained over time (Fig. 3F, 3hours), demonstrating that several enhancer elements co-locate within the same cell to stabilize folding of the domain. The angle around the TSS was reduced from ⁓100° to ⁓60° after E2 addition (Fig. 3G) further confirming folding back of the distal upstream region over the gene body. Concomitantly, PGR-TSS (Fos3) freedom of movement was increased 2-fold, from r=85 nm to r=172 nm, observed 3h post-stimulation (Fig. 3F).

In MCF7 cells, these results were coherent with the notion that folding is highly dynamic and variable from cell to cell (Cheng et al., 2020). Amplitude of variation in 3D positions appeared to be reduced as folding becomes stabilized when gene expression is stimulated by E2.

### Preexisting 3D structure of the PGR gene domain and regulatory region reorganizes during transcription activation

To further explore the structural properties of the PGR domain, we generated 3D models based on the spatial constraints measured by 5C contact frequencies using TADbit (Serra et al., 2017)(Fig. 4A). Specifically, each region of interest was represented as a chain of spherical beads each spanning 50 nm in diameter and containing 5 kb of DNA. In MCF7 cells, representations of an overlay ~10,000 models of the 500 kb active domain of PGR and enhancer region revealed a “croissant” shaped volume (Fig. 4A, domains shaded purple to pink). The adjacent TPCR6 gene formed a globular structure separated physically from the PGR domain as expected from seeing two distinct TADs in the 5C matrices in MCF7 cells (Fig.4A, red colored domain). Restraint-based modeling enabled extrapolating distances (d) between chosen fragments (Fig 4A). Grey arches in figure 4B are drawn between fragments which were separated by less than 50 nm in at least 50% of the calculated models of the PGR domain in MCF7 cells grown in hormone-starved medium (-E2) for three days. Genomic sites for which distances were shortened in at least 50% of E2 treated cells (+E2) are linked by red arches (Fig. 4B). This finding was coherent with a reduction in the 3D distances measured by 3D DNA FISH (Fig. 3C). Distances deduced from 5C-based models and three-way 3D FISH fell within the same distribution (Fig. 2C, 2D, 3E and 4B; Fig. Sup 3A) although the models tended toward smaller values. Obtaining the proper scale of models is not trivial for restraint-based modeling using 3C datasets (Trussart et al., 2015). The results confirm that the pre-existing PGR domain structure was reorganized upon estradiol treatment. In particular, numerous enhanced contacts demonstrate that RNA Pol2 bound upstream ERBS sites fold toward the gene body after 3h E2 exposure. Enhanced contacts are reminiscent of shortened distances measured in 3D FISH and the reduced dynamics of the loci relative to each other seen by ‘3 loci’ analysis (Fig. 3E-3G). Overall, the two arms of the modeled ‘croissant’ aligned and folded upon themselves (Fig. 4A). Strikingly, the adjacent TPCR6 TAD folding was not remodeled by estradiol. As expected from 5C data (Fig. 1), PGR domain folding and hormone induced reorganization was specific to MCF7 cells. Indeed, models from 5C data in MDA-MB-231 cells yielded distance distributions reminiscent a closed chromatin conformation of silent genes and, as expected, was not significantly altered following exposure to estradiol (Fig. S3B).

**Figure 4:**
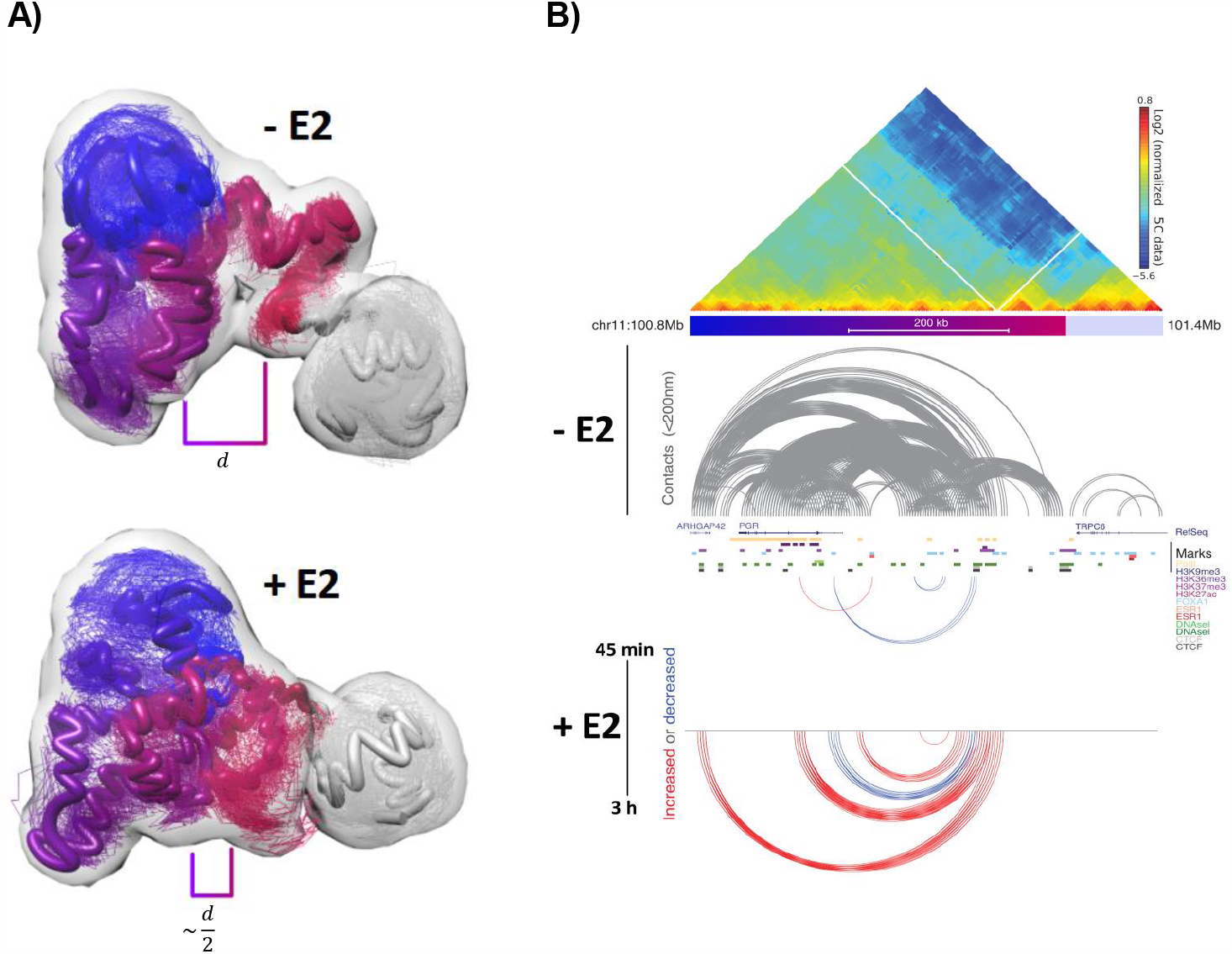
3D folding of the PGR chromatin domain reorganizes in response to transcription activation. A) 3D models 5C contact frequencies in MCF7 cells exposed to (+E2) or not (-E2) estradiol, 20% of the most frequent models using TadBit (Serra et al., 2017) with distances (d) between the PGR gene body and its upstream ERE region are displayed. B) 5C map (normalized counts) of the PGR domain on Chr 11. Color bar below the map corresponds to domains in A). Distances derived from models shown in A): grey arches link segments <200 nm in the maps, red and blue arches link segments for which distances increased (red) or decreased (blue) in at least 50% of the cell population after 45 min or 3h of estradiol exposure.

### Progressive ERα binding to enhancers mediates domain folding and transcription of PGR

We next investigated how transcription factors and cofactors operate within this 3D landscape. In MCF7 cells, expression of the PGR gene can be induced within minutes by adding E2. The ensuing mRNA synthesis is known to increase over time (Dalvai and Bystricky, 2010; Guertin et al., 2014;Shang et al., 2000) suggesting multiple steps of regulation. E2 binding to the ERα triggers a conformational change of the receptor enabling it to bind to its cognate binding sequences (ERB). Figure 5A shows progressive enrichment of ERα on all seven ERBSs within the PGR domain. Several chromatin-associated activators (GATA3, c-Fos, c-Jun, MYC, FoxA1), necessary for ERα-dependent gene activation were also recruited to the enhancers (Fig. 5B). In contrast, ERα did not associate with the TSS of PGR at any of the tested time-points prior to and following hormone addition. However, even in hormone starved cells, the TSS was H3K27 acetylated and bound by MYC (Fig. 5B) suggesting that the TSS is primed for transcription activation in MCF7 cells. Addition of E2 rapidly led to accumulation of MYC and of RNA Pol2 at the TSS. The binding profile of RNA Pol2 at the TSS and the PGR gene body varied over time. We thus calculated the RNA Pol2 pausing index and quantified mRNA synthesis using RNAseq and GROseq datasets (Fig. 5C). We observed a 1.5-fold (GROseq data) and a 6-fold (RNAseq data) increase of RNA within the early response (45 min) of E2 stimulation compared to the control situation. The RNA Pol2 peak declined rapidly from 5 min to 40 min (+E2), a time window during which PGR was modestly transcribed (Fig. 5C, early response). From 40 min to 160 min after E2 addition to the cells (late E2-response), PGR mRNA accumulation increased 4.5 times and RNA Pol2 was released from the TSS (Fig. 5C-late response). Finally, PGR expression levels stabilized until 160 min post induction concomitant with reduced RNA Pol2 pausing (Honkela et al., 2015; Shang et al., 2000).

**Figure 5:**
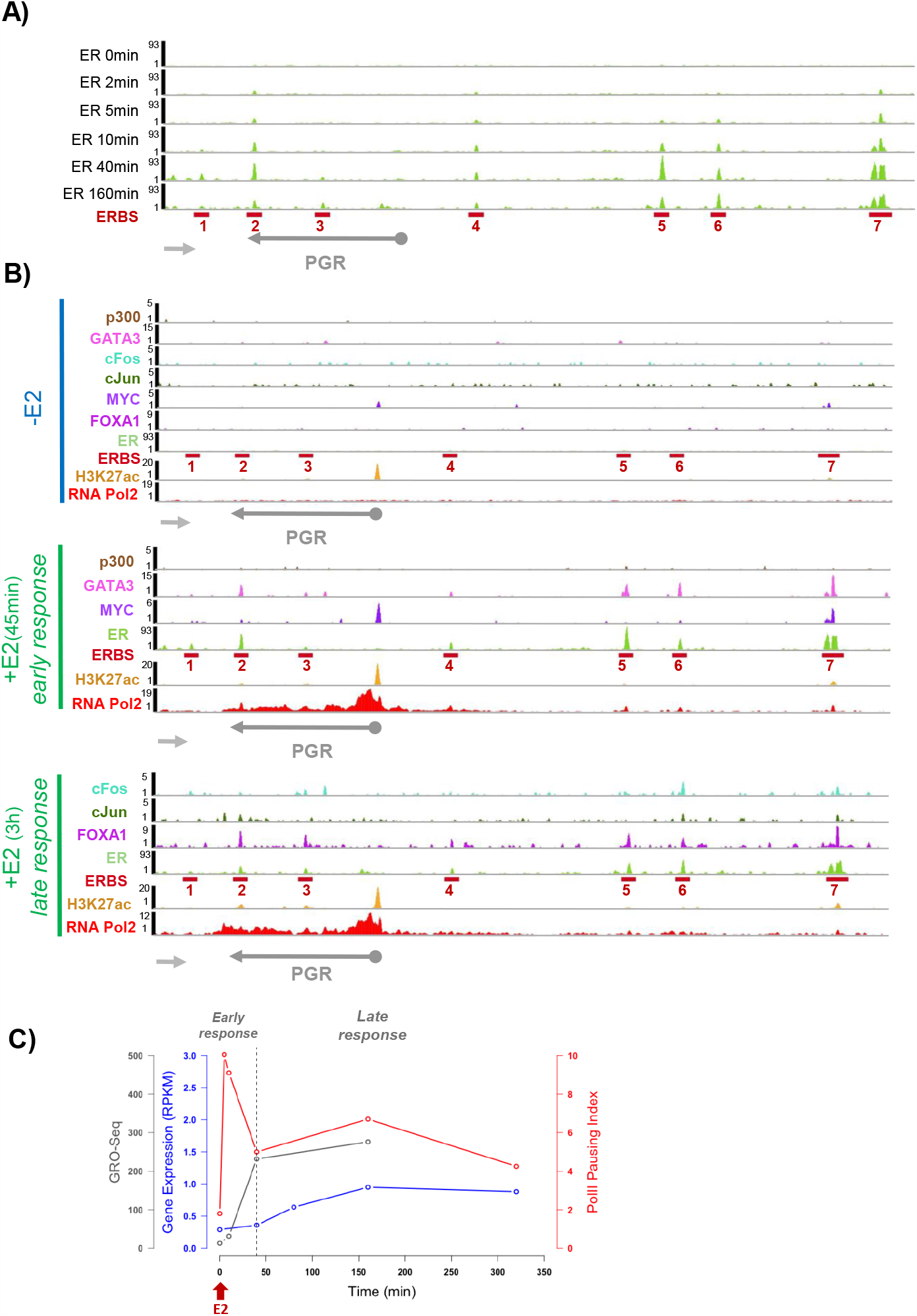
Domain folding facilitates enhancer function within the PGR chromatin domain during estradiol induced RNA polymerase 2 releases and activation. A) Kinetics of ERα progressive binding on ERBSs within regulatory domain of the PGR. Data analysed from (Guertin et al., 2014). B) Time-course of chromatin-associated activators (GATA3, P300, c-Fos, c-jun, MYC, FoxA1), necessary for ERa-dependent gene activation showing their presence at early or late E2-response. C) Time-course following estradiol addition to hormone starved MCF7 cells indicating RNA Pol 2 pausing inde6x (in red), mRNA production measured by RNA-seq (in blue) and by GRO-seq (in grey).

We propose that recruiting ERα to enhancers may enable enhancer interactions and conformational changes of the domain during steady state activation without direct enhancer-promoter contact. ERα accumulation thus correlates with RNA Pol2 release from a paused state and increased mRNA production (Fig. 5A, 5C and Fig. 6). ER-bound distal enhancer activity and chromatin looping hence fine-tune transcriptional output and permit discriminating transient responses from long-term sustained activity (Fig.6).

**Figure 6:**
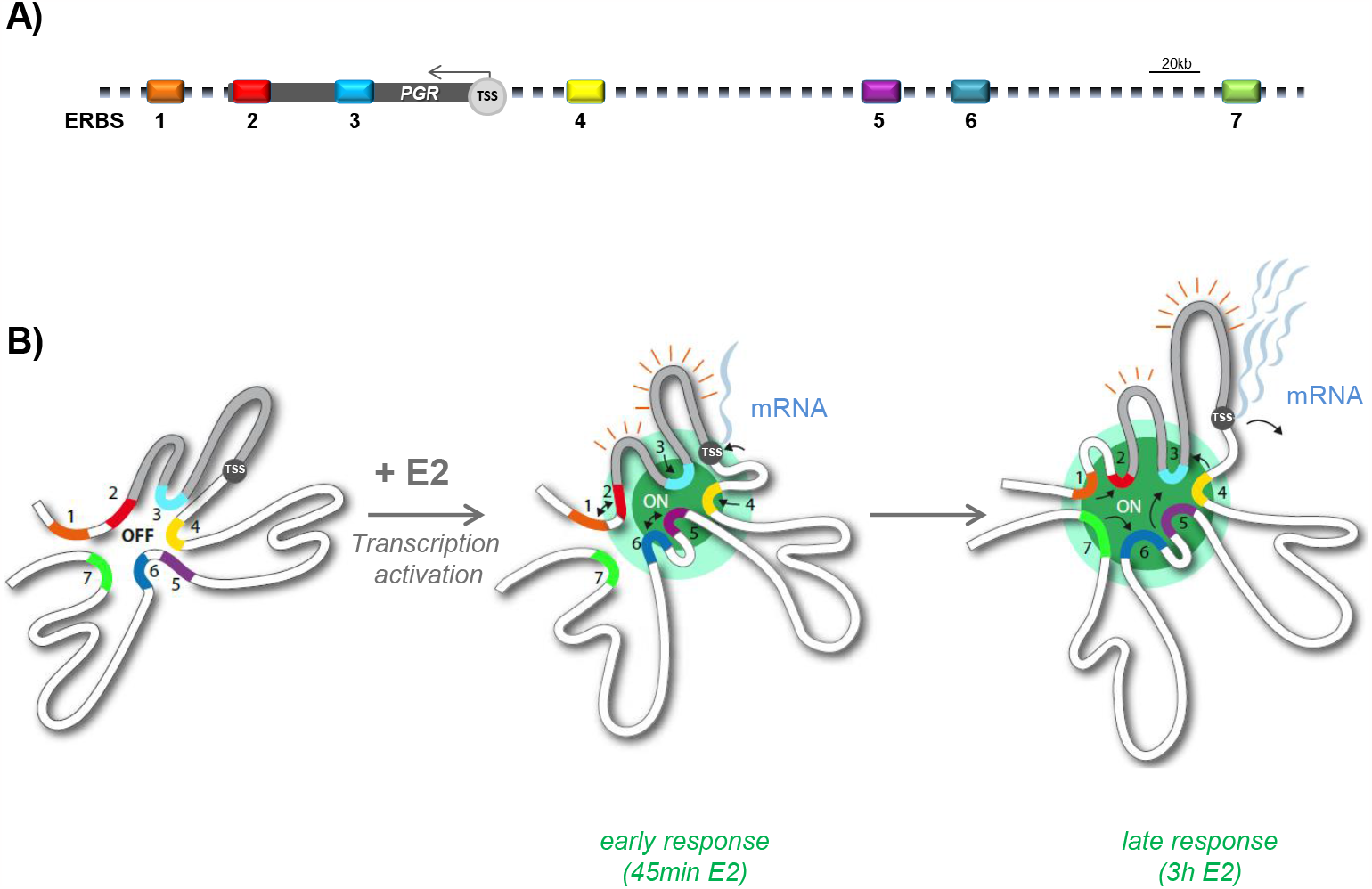
Three-dimensional PGR domain folding steps. A) Schematic cartoon showing linear region over 500 kb with the PGR gene body (100 kb), TSS and seven ER binding sites. B) Model of two-state conformations of the self-interacting PGR domain showing the representative contacts between proximal and distal enhancers (ERBSs) and TSS upon early and late estrogen response in MCF7 cells.

## Discussion

Here we show that chromatin fiber folding reflects transcriptional activity at the level of genomic domains and single genes. We discovered that domains encompassing silent genes show disordered structures that lack domain boundaries in triple negative human breast cancer cell lines reminiscent of silent domains in multiple cell lines (Cheng et al., 2020). In ERα-positive MCF7 cells, the same gene domains, namely PGR and ESR1, display TADs containing the gene body and its enhancer-spiked regulatory region, even in the absence of transcription. This pre-existing 3D structure is internally reorganized in response to hormone-induced transcription activation without altering the TAD borders. Reorganization appears to lock in already existing interactions rather than creating new ones, a phenomenon we could qualify as ‘caging in’. In particular at the PGR gene, the domain including the gene and the enhancer region is rapidly caged in. Caging is coherent with constrained motion of an E2 induced gene (Germier et al., 2017) and more generally with transcription induced reduced chromatin dynamics (Di Stefano et al., 2020; Nagashima et al., 2019; Shaban et al., 2020) (our unpublished observations in MCF7 cells). Concomitantly to caging, ERα accumulates at numerous enhancers within the region within less than one hour. These enhancers being close in space, ERα and associated transcription factors and cofactors create hubs. These hubs likely correspond to ERα foci seen across the nucleus in response to E2 stimulation of hundreds of genes (Kocanova et al., 2010b). Similar to ERα, progressive binding of the glucocorticoid receptor to distal enhancers within gene domains was proposed to lead to structural reorganization (Stavreva et al., 2015). Ligand bound nuclear receptors hence seem to induce folding between cis-acting enhancers and the adjacent gene domain. The resulting productive transcriptional conformation is similar to sustained active chromatin 3D hubs formed by the locus control region (LCR) of the globin genes in erythroid cells (Kooren et al., 2007) and by Sox2 (Stadhouders et al., 2017) but here as a mechanism to reversibly regulate acute activation.

Accumulation of ERα to enhancers may enable enhancer interactions and conformational changes of the domain during steady state activation (Fig. 3, 4). ERα accumulates progressively to reach a maximum concomitantly to the time of RNA Pol2 pause release and increased mRNA production (Fig. 5A). Hence, ER-bound distal enhancer activity and chromatin looping fine-tunes transcriptional output.

We found that ERα exclusively binds to non-promoter sequences of the PGR gene domain and not to the promoter of the PGR gene, an observation which challenges the common view that ERα is first recruited to the promoter of target genes where it triggers recruitment of cofactors and RNA Pol2. Genome-wide binding of ERα to non-promoter sequences was reported many years ago (Carroll et al., 2006) but its role was thus far not fully appreciated. In fact, numerous other transcription factors associate with distal regulatory elements rather than promoters directly (Shlyueva et al., 2014). It appears that multistep transcription factor binding to multiple enhancers and recruitment of co-factors creates a highly sensitive and reactive system to regulate transcription.

In addition to ERα, transcription factors including GATA3 and MYC were also recruited to enhancers within the PGR domain. GATA3 is a zinc-finger containing transcription factor important for cell differentiation and de-differentiation, in particular during EMT-like transition in metastatic breast cancer (Theodorou et al., 2013). GATA3 cooperates with ERα and is required to render cis-acting enhancers accessible for ER-mediated transcription activation (Tanaka et al., 2020; Theodorou et al., 2013). GATA3 binding was proposed to establish chromatin looping prior to activation (Theodorou et al., 2013) consistent with our observations that the PGR domain topology reflects transcriptional competence in MCF7 cells but not in MDA-MB-231 cells, which do not express GATA3. Moreover, GATA3 mutants were seen to disrupt regulatory networks enabling ER-mediated transcriptional response (Takaku et al., 2018) and repression of PGR.

Our study shows that 3D folding dynamics can be assessed using the 3-loci method to determine survival zones of linked genomic loci as a direct approach to analyze stimulus induced changes of specific domains without the use of sequencing-based chromosome conformation capture methodologies. We have demonstrated here that 3D models based on chromosome conformation capture data confirm the picture drawn from quantitative 3D imaging of specific loci. The 3-loci method is applicable at any scale and to any cell type.

We propose that recruiting ERα to enhancers may enable enhancer interactions and topological changes of the domain during steady state activation without direct enhancer-promoter contact. ER-bound distal enhancer activity and chromatin looping fine-tune transcriptional output and may permit discriminating transient responses from sustained activity (Fig.6). Preestablished chromatin architectures control gene expression without the need for de-novo long range rewiring of contacts. In fact, the common breast cancer cell lines used here may represent states of genome adaptation to optimize proliferation and response to physiological environments. An attractive hypothesis could thus be that selective estrogen receptor modulator anti-estrogens exploit such environment to hijack preset gene regulatory domains. It may therefore be relevant to explore, and possibly act upon, 3D domain organization when therapeutic resistance or recurrence appears (Fukuoka et al., 2022).

## Material and methods

### Cell culture

The human ERα-positive breast cancer cell line MCF7 and the ERα−negative breast cancer cell line MDA-MB-231 were purchased from ATTC and maintained in DMEM F-12 (Gibco) for MCF7 or DMEM (Gibco) for MDA-MB231 with Glutamax and complemented with 50μg/ml gentamicin, 1mM sodium pyruvate and 10% FCS. Cells were grown at 37°C in a humidified atmosphere containing 5% CO_2_.

To study the effects of 17β-estradiol (E2) on oestrogen-regulated genes (Progesterone receptor gene (PGR), Estrogen receptor gene (ESR1), Cyclin D1 gene (CCND1) and GREB1) cells were grown for 3 days in phenol red-free media supplemented with 10% charcoal-stripped FCS (-E2) and subsequently treated with 100nM E2 (Sigma) for the indicated times.

### DNA-FISH

3D DNA FISH experiments were performed as previously described in (Kocanova et al., 2018, 2010a). Briefly, cells were grown for 3 days on 10mm round glass coverslips in 24-well plates using DMEM (DMEM/F-12 for MCF7 cells) without phenol red, containing 10% charcoal-stripped FCS, before addition of 100nM E2 for the indicated times. Coverslips were then washed once with PBS, fixed in freshly-made 4% paraformaldehyde (pFA)/PBS for 10 min at room temperature and during the last three minutes a 200μl of 0.5% Triton X-100/PBS were added homogenously. From this step, the cells were treated as followed, at room temperature and with moderate shaking. Cells were washed three-times for 3 min in 0.01% Triton X-100/PBS, incubated in 0.5% Triton X-100/PBS for 10 min at room temperature and treated or not with 0.2 mg/ml RNase A in 2xSSC for 30 min at 37°C. After 3 washes of 10 min in PBS, cells were incubated in 0.1M HCl for 5 min (for MCF7 cells) or in 0.05M HCl for 2 min (for MDA-MB231cells), washed twice in 2xSSC for 3 min and then left in 50% formamide/2xSSC (pH=7.2) 1h minimum at room temperature or one week maximum at 4°C before being used for 3D DNA FISH.

Fosmids were purchased at C.H.O.R.I. (see Suppl. Table 2) and were labeled using the BioPrime DNA Labeling System from Invitrogen by incorporation of fluorochrome-conjugated nucleotides Atto647N-dUTP-NT (Jena Bioscience), atto550-dUTP-NT (Jean Biosciences) or ChromaTide® AlexaFluor® 488-5-dUTP (Molecular Probe). Labeled probes were then purified trough a G50 column from the Illustra MicroSpin G50 kit (GE Healthcare) and precipitated overnight with Salmon sperm DNA (Sigma), Human Cot-1 DNA (Invitrogen), 3M NaAc and 100% cold EtOH. After centrifugation, the pellet was resuspended in an appropriate volume of HP (Hybridization Premix, containing 50% formamide in a buffer solution) to obtain equal concentration (100 ng/µl) for all fosmid probes. For DNA-FISH, 200ng of labeled fosmids were added to the prepared coverslips. The coverslips were sealed with rubber cement (Electron Microscopy Sciences) and placed into a hybridizer (Dako). Denaturation of the probes and target DNA was performed simultaneously at 85°C for 2 minutes and samples were then incubated overnight at 37°C. Before microscopy, coverslips were then washed, with gentle shaking, 4 times for 3 min in 2xSSC at 45°C and 4 times for 3 min in 0.1x SSC at 60°C before being mounted with 4μl of Vectashield® Antifade Mounting Medium with DAPI at 1.5μg/ml and sealed with transparent nail polish.

### Image acquisition and analysis

3D-DNA FISH observations were performed using an Olympus IX-81 wide-field fluorescence microscope, equipped with a CoolSNAP HQ CCD camera (Photometrics), a Polychrome V monochromator (Till Photonics) equipped with a 150 W xenon source used with a 15 nm bandwidth, an electric PIFOC piezo stepper (PI) with an accuracy of 10 nm, and imaged through an Olympus oil immersion objective 100x PLANAPO NA1.4. Acquisitions of DAPI, Alexa 488, ATTO550 and ATTO647 fluorophores were performed using multiband dichroic mirrors (Chroma), specific single-band emission filters mounted on a motorized wheel (PI), and emission filters ET450/40, ET520/40, ET580/40, and ET685/60. Image acquisitions were performed in ~21 focal planes with 200 nm step size for 3D DNA FISH. This configuration was driven by MetaMorph**®** (Microscopy Automation and Image Analysis Software from Molecular Devices). Images were analysed using a home-made script running on ImageJ (Kocanova et al., 2018). Inter-probe distances were determined from 80-130 nuclei for each experimental condition and the significance of any difference in the data distributions was assessed using Fisher’s test. A p-value ≤0.05 was considered statistically significant.

### ChIP-seq processing

Published ChIPseq data in MCF7 were downloaded from GEO DataSets (https://www.ncbi.nlm.nih.gov/geo/) and treated as subsequently described. Raw data were downloaded with fastq-dump from sratoolkit (2.8.2-1). (http://ncbi.github.io/sra-tools/). The quality of the reads was estimated with FastQC (0.11.7). Sequencing reads were aligned to the reference human genome assembly (hg19/GRCh37) using Burrows-Wheeler Aligner (Honkela et al., 2015; Li and Durbin, 2010) 2010) (0.7.17) with default parameters. ER peak calling was performed using MACS2 (Zhang et al., 2008). ERBS were defined according to ER peak calling in +E2 40min condition. Bigwig files were generated and normalized (RPKM) using scripts from deepTools utilities. (3.0.2) (Ramírez et al., 2016). Paused RNA Pol2 indices were defined as the ratio of RNA Pol2 (total) density in the promoter-proximal region (-30 bp to TSS from + 300 bp) to the total RNA Pol2 density in the transcribed regions (TSS + 300 bp to TES). 5C matrices were visualized using WashU Epigenome Browser(Li et al., 2019). ER cofactors and epigenetic marks were defined and analyzed according to published data: Supplementary Table 1.

### RNA-seq processing

RNA-Seq time course in MCF7 before and after E2 treatment were downloaded from GEO (accession number: GSE62789) (Honkela et al., 2015). Raw data were downloaded with fastq-dump from sratoolkit (2.8.2-1). The quality of the reads was estimated with FastQC (0.11.7). Alignment was performed using STAR (2.6.0c) to the hg19/GRCh37 reference human genome assembly. Gene expression values were quantified from RNA sequencing data using HTSeq (0.9.1) and were then normalized (RPKM) in both cell lines.

### 5C primer design

5C primers were designed using My5C (Lajoie et al. Nature Methods 2009). 5C employs two types of primers: 5C forward and 5C reverse primers. We used an alternating primer where a single 5C primer was designed for each HindIII fragment throughout the genomic regions analyzed here, so that forward and reverse primers alternate (see (Dostie et al., 2006)).

### 5C data processing

We used My5C (Lajoie et al., 2009) to design 5C primers, using default settings. We designed 5F forward and reverse primers using the alternating primer design(Dostie et al., 2006). 5C data analysis consist of alignment, noise removal, scaling, binning, and balancing (iterative correction). First, we aligned the 5C sequencing reads to the reference primer set using Novoalign (version 3.02.00) to determine the interactions between primer pairs. Any primer pair or individual primer that has very low or excessive number of interactions will be removed from the matrix using z-score in both cis and trans using thresholds 6 and 12 respectively. Then the matrix was be read-normalized to adjust for the number of sequencing reads per sample. Finally, we binned and balanced the matrix to decrease bias, complexity, and get multi resolution data using the ICE method (Imakaev et al. (2012)): Data was binned in 15 kb bins with 8 steps which creates 13.125 kb overlaps between bins. Balancing uses Sinkhorn-Knopp algorithm which rescales the rows and columns by dividing their sums by their means iteratively to get matrix convergence. Thus, the total number of interactions per primer (locus) should be the same. LOWESS method (Locally Weighted Regression: An Approach to Regression Analysis by Local Fitting) was used to estimate the expected interactions for given distances.

### 5C data analysis

Interaction counts by genomic distance for estrogen-regulated gene regions (PGR, ESR1, CCND1 and GREB1) have been computed with hicPlotDistVsCounts tool from HiCExplorer (2.2.1.1) utilities. Correlation Heatmaps were computed with hicCorrelate tool from HiCExplorer. In order to study Interaction counts distributions at ERBS/ERBS and ERBS/TSS for the PGR gene, we considered a window region from each ER peak summit +/- 10kb (+/- 15kb for ERE7) within the PGR gene region matrices and PGR gene TSS +/- 10kb. We then plotted the corresponding averaged interaction count values around the two interacting regions, and compared each of the two E2 induction time-point distributions. Violin Plots have been generated using R home-made scripts.

### Modeling

5C produces two-dimensional matrices that represent the frequency of interactions between loci along the genomic region of interest. To transform such data into a 3D conformation of higher-order chromatin folding, we used TADbit (Serra et al., 2017). Structure determination by TADbit can be seen as an iterative series of three main steps: translating the data into spatial restraints, constructing an ensemble of structures that satisfy these restraints, and analyzing the ensemble to produce the final structure. Specifically, each particle pair in the models was restrained by a series of harmonic oscillator centered on a distance derived from the 5C data as previously described (Dostie et al., 2006; Serra et al., 2017). A total of 1,000 models at 5Kb resolution per region were built by TADbit with *maxdist=250, upfreq=0, lowfreq=-0*.*6, scle=0*.*01* as input parameters. The Pearson correlation coefficient between a contact map obtained from the optimal models and the input 5C matrix was 0.94, indicative of accurate models (Trussart et al., 2015). All remining TADbit *model* parameters were set to default values. The resulting models were further analyzed to obtain contact arches analysis, distance distributions as well as visual representations of the models.

*Contact arches*: A “contact arch” was defined between two model particles if their distance was less than 200nm in at least 50% of the models in the ensemble. To assess the number of increased/decreased contacts, we computed the difference in models having such arch. An increase of contacts in at least 20% of models with respect -E2 models indicated an increase of an arch (red color). Conversely, a decrease of contacts in at least 20% of models with respect -E2 models indicated a decrease of an arch (blue color).

*Distance analysis:* Euclidean distances in Cartesian space between selected fosmids were calculated for the entire ensemble and represented as box plots. If a fosmid occupied more than one particle, the center of mass of the constituting particles was used as the *x,y,z* coordinates of the fosmid.

*Ensemble visualization*: the UCSF Chimera package (Yang et al., 2012), a highly extensible program for interactive visualization of molecular structures, was used to produce all images of the models. Models were superimposed and visualized with a wire representation within a transparent molecular density map of the occupancy of all particles in the models (obtained using the *molmap* function in Chimera). Finally, the “centroid” model (that is, the model central to the superimposed ensemble) was represented as a thick worm like structure for easy visualization (obtained using the *shape tube* function in Chimera).

### Computing survival radii – 3 loci analysis

Distances between the three loci R, G, B were measured in 3D (see above).

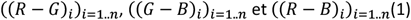

In the plane defined by the three loci, we denote by *d*_*rgmin*_, *d*_*rbmin*_, *d*_*gbmin*_the minimal distances) and *d*_*rgmax*_, *d*_*rbmax*_, *d*_*gbmax*_the maximal distances between R-G-B, respectively

We can note that

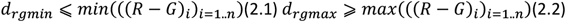

We have the same inequalities for couples (*d*_*rbmin*_, *d*_*rbmax*_)and (*d*_*gbmin*_, *d*_*gbmax*_).

In the limit case we make the assumption that when ***n*** is large enough:

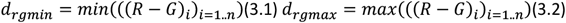

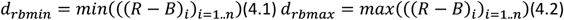

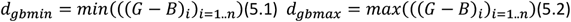

We define the following relations :

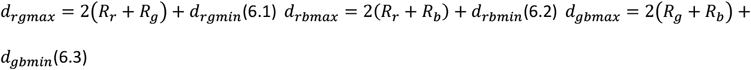

Solving this equation gives

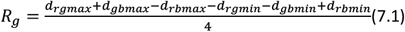

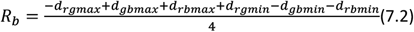

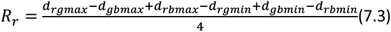

When applying the following change of variable :

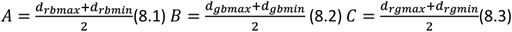

we can specify the position of the third locus in relation to the other loci by

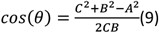

Where θ is the angle between Red-Green and Green-Blue axes.

For each triplet of measured distances, ((*R* − *G*)_*i*_, (*G* − *B*)_*i*_, (*R* − *B*)_*i*_)_*i*=1..*n*_the positions of the loci

((*x*_*r*_, *y*_*r*_), (*x*_*g*_, *y*_*g*_), (*x*_*b*_, *y*_*b*_)) were deduced as

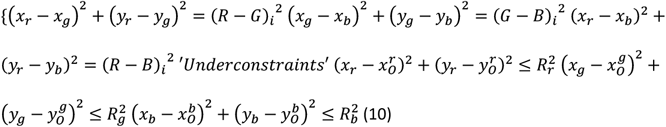

To solve system (10) the three survival zones using polar coordinates were calculated:

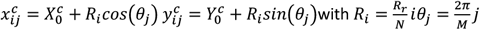

where *c* is the color index **R, G** or **B**. *N* is the number of survival zones subdivision and M the polar angle subdivision.

## Supporting information

Image File-supl

## Code and data availability

Code and data can be accessed at https://github.com/FlavienRaynal/PGR_Kocanova for 5C and

ChIPseq analyses and at https://src.koda.cnrs.fr/alain.kamgoue.3/tripleloci for triple loci.

## Acknowledgements

We acknowledge support from the light imaging Toulouse CBI facility (LITC).

This work was supported by grants from HFSPO RGP0044 to JD, MMR, KB; Agence Nationale de la Recherche (ANR): SINFONIE AAPG-18-CE12-006 to KB. MAM-R acknowledges support by the Spanish Ministerio de Ciencia e Innovación (PID2020-115696RB-I00). JD acknowledges support from the National Human Genome Research Institute (HG003143). JD is an investigator of the Howard Hughes Medical Institute.

## Author contributions

MMR, JD, KB study conception and coordination, data analysis, manuscript drafting

Silvia Kocanova FISH experiments, image acquisition, data analysis, manuscript

Flavien Raynal bioinformatic analyses

Isabelle Goiffon FISH experiments

Betul Agkol Oksuz 5C analysis

Davide Baú 3D modeling

Alain Kamgoué mathematics

Sylvain Cantaloube image analysis

Ye Zhan 5C experiments

Bryan Lajoie 5C primer design

## References

Abbas, A., He, X., Niu, J., Zhou, B., Zhu, G., Ma, T., Song, J., Gao, J., Zhang, M.Q., Zeng, J., 2019. Integrating Hi-C and FISH data for modeling of the 3D organization of chromosomes. Nat Commun 10, 2049. 10.1038/s41467-019-10005-6

Abdalla, M.O.A., Yamamoto, T., Maehara, K., Nogami, J., Ohkawa, Y., Miura, H., Poonperm, R., Hiratani, I., Nakayama, H., Nakao, M., Saitoh, N., 2019. The Eleanor ncRNAs activate the topological domain of the ESR1 locus to balance against apoptosis. Nat Commun 10, 3778. 10.1038/s41467-019-11378-4

Boninsegna, L., Yildirim, A., Polles, G., Zhan, Y., Quinodoz, S.A., Finn, E.H., Guttman, M., Zhou, X.J., Alber, F., 2022. Integrative genome modeling platform reveals essentiality of rare contact events in 3D genome organizations. Nat Methods 19, 938–949. 10.1038/s41592-022-01527-x

Carroll, J.S., Meyer, C. a, Song, J., Li, W., Geistlinger, T.R., Eeckhoute, J., Brodsky, A.S., Keeton, E.K., Fertuck, K.C., Hall, G.F., Wang, Q., Bekiranov, S., Sementchenko, V., Fox, E. a, Silver, P. a, Gingeras, T.R., Liu, X.S., Brown, M., 2006. Genome-wide analysis of estrogen receptor binding sites. Nature genetics 38, 1289–97. 10.1038/ng1901

Cheng, R.R., Contessoto, V.G., Lieberman Aiden, E., Wolynes, P.G., Di Pierro, M., Onuchic, J.N., 2020. Exploring chromosomal structural heterogeneity across multiple cell lines. eLife 9, e60312. 10.7554/eLife.60312

Dalvai, M., Bystricky, K., 2010. Cell Cycle and Anti-Estrogen Effects Synergize to Regulate Cell Proliferation and ER Target Gene Expression. PLOS ONE 5. 10.1371/journal.pone.0011011

Di Stefano, M., Stadhouders, R., Farabella, I., Castillo, D., Serra, F., Graf, T., Marti-Renom, M.A., 2020. Transcriptional activation during cell reprogramming correlates with the formation of 3D open chromatin hubs. Nat Commun 11, 2564. 10.1038/s41467-020-16396-1

Dostie, J., Richmond, T.A., Arnaout, R.A., Selzer, R.R., Lee, W.L., Honan, T.A., Rubio, E.D., Krumm, A., Lamb, J., Nusbaum, C., Green, R.D., Dekker, J., 2006. Chromosome Conformation Capture Carbon Copy (5C): A massively parallel solution for mapping interactions between genomic elements. Genome Res. 16, 1299–1309. 10.1101/gr.5571506

ENCODE Project Consortium, 2012. An integrated encyclopedia of DNA elements in the human genome. Nature 489, 57–74. 10.1038/nature11247

Fukuoka, M., Ichikawa, Y., Osako, T., Fujita, T., Baba, S., Takeuchi, K., Tsunoda, N., Ebata, T., Ueno, T., Ohno, S., Saitoh, N., 2022. The ELEANOR noncoding RNA expression contributes to cancer dormancy and predicts late recurrence of estrogen receptor‐positive breast cancer. Cancer Science 113, 2336–2351. 10.1111/cas.15373

Fullwood, M.J., Liu, M.H., Pan, Y.F., Liu, J., Xu, H., Mohamed, Y.B., Orlov, Y.L., Velkov, S., Ho, A., Mei, P.H., Chew, E.G.Y., Huang, P.Y.H., Welboren, W.-J., Han, Y., Ooi, H.S., Ariyaratne, P.N., Vega, V.B., Luo, Y., Tan, P.Y., Choy, P.Y., Wansa, K.D.S.A., Zhao, B., Lim, K.S., Leow, S.C., Yow, J.S.,Joseph, R., Li, H., Desai, K.V., Thomsen, J.S., Lee, Y.K., Karuturi, R.K.M., Herve, T., Bourque, G., Stunnenberg, H.G., Ruan, X., Cacheux-Rataboul, V., Sung, W.-K., Liu, E.T., Wei, C.-L., Cheung, E., Ruan, Y., 2009. An oestrogen-receptor-α-bound human chromatin interactome. Nature 462, 58–64. 10.1038/nature08497

Germier, T., Kocanova, S., Walther, N., Bancaud, A., Shaban, H.A., Sellou, H., Politi, A.Z., Ellenberg, J., Gallardo, F., Bystricky, K., 2017. Real-time chromatin dynamics at the single gene level during transcription activation. bioRxiv 111179. 10.1101/111179

Gheldof, N., Smith, E.M., Tabuchi, T.M., Koch, C.M., Dunham, I., Stamatoyannopoulos, J.A., Dekker, J., 2010. Cell-type-specific long-range looping interactions identify distant regulatory elements of the CFTR gene. Nucleic Acids Res 38, 4325–4336. 10.1093/nar/gkq175

Giamarchi, C., Solanas, M., Chailleux, C., Augereau, P., Vignon, F., Rochefort, H., Richard-Foy, H., 1999. Chromatin structure of the regulatory regions of pS2 and cathepsin D genes in hormone-dependent and -independent breast cancer cell lines. Oncogene. 10.1038/sj.onc.1202317

Gibcus, J.H., Dekker, J., 2013. The Hierarchy of the 3D Genome. Molecular Cell 49, 773–782. 10.1016/j.molcel.2013.02.011

Giorgetti, L., Heard, E., 2016. Closing the loop: 3C versus DNA FISH. Genome Biol 17, 215. 10.1186/s13059-016-1081-2

Guertin, M.J., Zhang, X., Coonrod, S.A., Hager, G.L., 2014. Transient Estrogen Receptor Binding and p300 Redistribution Support a Squelching Mechanism for Estradiol-Repressed Genes. Molecular Endocrinology 28, 1522–1533. 10.1210/me.2014-1130

Honkela, A., Peltonen, J., Topa, H., Charapitsa, I., Matarese, F., Grote, K., Stunnenberg, H.G., Reid, G., Lawrence, N.D., Rattray, M., 2015. Genome-wide modeling of transcription kinetics reveals patterns of RNA production delays. Proceedings of the National Academy of Sciences of the United States of America. 10.1073/pnas.1420404112

Jardin, I., Diez-Bello, R., Lopez, J.J., Redondo, P.C., Salido, G.M., Smani, T., Rosado, J.A., 2018. Trpc6 channels are required for proliferation, migration and invasion of breast cancer cell lines by modulation of orai1 and orai3 surface exposure. Cancers. 10.3390/cancers10090331

Kim, S., Shendure, J., 2019. Mechanisms of Interplay between Transcription Factors and the 3D Genome. Molecular Cell 76, 306–319. 10.1016/j.molcel.2019.08.010

Kim, T.H., Dekker, J., 2018. 5C Analysis of 3C, ChIP-Loop, and Control Libraries. Cold Spring Harb Protoc 2018, pdb.prot097899. 10.1101/pdb.prot097899

Kininis, M., Chen, B.S., Diehl, A.G., Isaacs, G.D., Zhang, T., Siepel, A.C., Clark, A.G., Kraus, W.L., 2007. Genomic Analyses of Transcription Factor Binding, Histone Acetylation, and Gene Expression Reveal Mechanistically Distinct Classes of Estrogen-Regulated Promoters. Mol Cell Biol 27, 5090–5104. 10.1128/MCB.00083-07

Kılıç, Y., Çelebiler, A.Ç., Sakızlı, M., 2014. Selecting housekeeping genes as references for the normalization of quantitative PCR data in breast cancer. Clin Transl Oncol 16, 184–190. 10.1007/s12094-013-1058-5

Kocanova, S., Goiffon, I., Bystricky, K., 2018. 3D FISH to analyse gene domain-specific chromatin remodeling in human cancer cell lines. Methods 142, 3–15. 10.1016/j.ymeth.2018.02.013

Kocanova, S., Kerr, E., Rafique, S., Boyle, S., Boyle, S., Katz, E., Caze-Subra, S., Bickmore, W.A., Bystricky, K., 2010a. Activation of Estrogen-Responsive Genes Does Not Require Their Nuclear Co-Localization. PLOS Genetics 6. 10.1371/journal.pgen.1000922

Kocanova, S., Mazaheri, M., Caze-Subra, S., Bystricky, K., 2010b. Ligands specify estrogen receptor alpha nuclear localization and degradation. BMC cell biology 11, 98.

Kooren, J., Palstra, R.-J., Klous, P., Splinter, E., von Lindern, M., Grosveld, F., de Laat, W., 2007. β-Globin Active Chromatin Hub Formation in Differentiating Erythroid Cells and in p45 NF-E2 Knock-out Mice. Journal of Biological Chemistry 282, 16544–16552. 10.1074/jbc.M701159200

Lajoie, B.R., van Berkum, N.L., Sanyal, A., Dekker, J., 2009. My5C: web tools for chromosome conformation capture studies. Nat Methods 6, 690–691. 10.1038/nmeth1009-690

Lassadi, I., Kamgoue, A., Goiffon, I., Tanguy-le-Gac, N., Bystricky, K., 2015. Differential chromosome conformations as hallmarks of cellular identity revealed by mathematical polymer modeling. PLOS Computational Biology 11. 10.1371/journal.pcbi.1004306

Le Dily, F.L., Bau, D., Pohl, A., Vicent, G.P., Serra, F., Soronellas, D., Castellano, G., Wright, R.H.G., Ballare, C., Filion, G., Marti-Renom, M.A., Beato, M., 2014. Distinct structural transitions of chromatin topological domains correlate with coordinated hormone-induced gene regulation. Genes and Development 28, 2151–2162. 10.1101/gad.241422.114

Li, D., Hsu, S., Purushotham, D., Sears, R.L., Wang, T., 2019. WashU Epigenome Browser update 2019. Nucleic Acids Research 47, W158–W165. 10.1093/nar/gkz348

Li, H., Durbin, R., 2010. Fast and accurate long-read alignment with Burrows–Wheeler transform. Bioinformatics 26, 589–595. 10.1093/bioinformatics/btp698

Luo, Y., Hitz, B.C., Gabdank, I., Hilton, J.A., Kagda, M.S., Lam, B., Myers, Z., Sud, P., Jou, J., Lin, K., Baymuradov, U.K., Graham, K., Litton, C., Miyasato, S.R., Strattan, J.S., Jolanki, O., Lee, J.-W., Tanaka, F.Y., Adenekan, P., O’Neill, E., Cherry, J.M., 2020. New developments on the Encyclopedia of DNA Elements (ENCODE) data portal. Nucleic Acids Res 48, D882–D889. 10.1093/nar/gkz1062

Nagashima, R., Hibino, K., Ashwin, S.S., Babokhov, M., Michael Babokhov, Fujishiro, S., Imai, R., Nozaki, T., Tamura, S., Tani, T., Kimura, H., Shribak, M., Kanemaki, M.T., Sasai, M., Maeshima, K., 2019. Single nucleosome imaging reveals loose genome chromatin networks via active RNA polymerase II. Journal of Cell Biology 218, 1511–1530. 10.1083/jcb.201811090

Nir, G., Farabella, I., Pérez Estrada, C., Ebeling, C.G., Beliveau, B.J., Sasaki, H.M., Lee, S.D., Nguyen, S.C., McCole, R.B., Chattoraj, S., Erceg, J., AlHaj Abed, J., Martins, N.M.C., Nguyen, H.Q.,Hannan, M.A., Russell, S., Durand, N.C., Rao, S.S.P., Kishi, J.Y., Soler-Vila, P., Di Pierro, M., Onuchic, J.N., Callahan, S.P., Schreiner, J.M., Stuckey, J.A., Yin, P., Aiden, E.L., Marti-Renom, M.A., Wu, C. -ting, 2018. Walking along chromosomes with super-resolution imaging, contact maps, and integrative modeling. PLoS Genet 14, e1007872. 10.1371/journal.pgen.1007872

Oudelaar, A.M., Beagrie, R.A., Kassouf, M.T., Higgs, D.R., 2021. The mouse alpha-globin cluster: a paradigm for studying genome regulation and organization. Curr Opin Genet Dev 67, 18–24. 10.1016/j.gde.2020.10.003

Paliou, C., Guckelberger, P., Schöpflin, R., Heinrich, V., Esposito, A., Chiariello, A.M., Bianco, S., Annunziatella, C., Helmuth, J., Haas, S., Jerković, I., Brieske, N., Wittler, L., Timmermann, B.,Nicodemi, M., Vingron, M., Mundlos, S., Andrey, G., 2019. Preformed chromatin topology assists transcriptional robustness of Shh during limb development. Proc. Natl. Acad. Sci.U.S.A. 116, 12390–12399. 10.1073/pnas.1900672116

Ramírez, F., Ryan, D.P., Grüning, B., Bhardwaj, V., Kilpert, F., Richter, A.S., Heyne, S., Dündar, F., Manke, T., 2016. deepTools2: a next generation web server for deep-sequencing data analysis. Nucleic Acids Res 44, W160–W165. 10.1093/nar/gkw257

Serra, F., Baù, D., Goodstadt, M., Castillo, D., Filion, G.J., Marti-Renom, M.A., 2017. Automatic analysis and 3D-modelling of Hi-C data using TADbit reveals structural features of the fly chromatin colors. PLoS Comput Biol 13, e1005665. 10.1371/journal.pcbi.1005665

Shaban, H.A., Barth, R., Roman Barth, Recoules, L., Bystricky, K., 2020. Hi-D: nanoscale mapping of nuclear dynamics in single living cells. Genome Biology 21, 95–95. 10.1186/s13059-020-02002-6

Shang, Y., Hu, X., DiRenzo, J., Lazar, M.A., Brown, M., 2000. Cofactor Dynamics and Sufficiency in Estrogen Receptor–Regulated Transcription. Cell 103, 843–852. 10.1016/S0092-8674(00)00188-4

Shlyueva, D., Stampfel, G., Stark, A., 2014. Transcriptional enhancers: From properties to genome-wide predictions. Nature Reviews Genetics. 10.1038/nrg3682

Sikorska, N., Sexton, T., 2020. Defining Functionally Relevant Spatial Chromatin Domains: It is a TAD Complicated. Journal of Molecular Biology 432, 653–664. 10.1016/j.jmb.2019.12.006

Smith, E.M., Lajoie, B.R., Jain, G., Dekker, J., 2016. Invariant TAD Boundaries Constrain Cell-Type-Specific Looping Interactions between Promoters and Distal Elements around the CFTR Locus. Am J Hum Genet 98, 185–201. 10.1016/j.ajhg.2015.12.002

Stadhouders, R., Vidal, E., Serra, F., Di Stefano, B., Le Dily, F., Quilez, J., Gomez, A., Collombet, S., Berenguer, C., Cuartero, Y., Hecht, J., Filion, G., Beato, M., Marti-Renom, M.A., Graf, T.,Stadhouders, R., Marc Marti-Renom, dr A., Thomas Graf, D., 2017. Transcription factors orchestrate dynamic interplay between genome topology and gene regulation during cell reprogramming. bioRxiv. 10.1101/132456

Stavreva, D. a, Coulon, A., Baek, S., Sung, M., John, S., Stixova, L., Tesikova, M., Hakim, O., Miranda, T., Hawkins, M., Stamatoyannopoulos, J. a, Chow, C.C., Hager, G.L., 2015. Dynamics of chromatin accessibility and long-range interactions in response to glucocorticoid pulsing. Genome Research 25, 845–857. 10.1101/gr.184168.114.8

Szabo, Q., Donjon, A., Jerković, I., Ivana Jerković, Papadopoulos, G.L., Cheutin, T., Bonev, B.B., Nora, E.P., Bruneau, B.G., Bantignies, F., Cavalli, G., 2020. Regulation of single-cell genome organization into TADs and chromatin nanodomains. Nature Genetics 52, 1151–1157. 10.1038/s41588-020-00716-8

Takaku, M., Grimm, S.A., Roberts, J.D., Chrysovergis, K., Bennett, B.D., Myers, P., Perera, L., Tucker, C.J., Perou, C.M., Wade, P.A., 2018. GATA3 zinc finger 2 mutations reprogram the breast cancer transcriptional network. Nature Communications 9, 1–14. 10.1038/s41467-018-03478-4

Tan, L., Ma, W., Wu, H., Zheng, Y., Xing, D., Chen, R., Li, X., Daley, N., Deisseroth, K., Xie, X.S., 2021. Changes in genome architecture and transcriptional dynamics progress independently of sensory experience during post-natal brain development. Cell 184, 741-758.e17. 10.1016/j.cell.2020.12.032

Tanaka, H., Takizawa, Y., Takaku, M., Grimm, S.A., Wade, P.A., Kurumizaka, H., 2020. GATA3 with nucleosomes. Nature Communications. 10.1038/s41467-020-17959-y

Theodorou, V., Stark, R., Menon, S., Carroll, J.S., 2013. GATA3 acts upstream of FOXA1 in mediating ESR1 binding by shaping enhancer accessibility. Genome Research 23, 12–22. 10.1101/gr.139469.112

Trussart, M., Serra, F., Baù, D., Junier, I., Serrano, L., Marti-Renom, M.A., 2015. Assessing the limits of restraint-based 3D modeling of genomes and genomic domains. Nucleic Acids Research 43, 3465–3477. 10.1093/nar/gkv221

van Steensel, B., Furlong, E.E.M., 2019. The role of transcription in shaping the spatial organization of the genome. Nat Rev Mol Cell Biol. 10.1038/s41580-019-0114-6

Yang, Z., Lasker, K., Schneidman-Duhovny, D., Webb, B., Huang, C.C., Pettersen, E.F., Goddard, T.D., Meng, E.C., Sali, A., Ferrin, T.E., 2012. UCSF Chimera, MODELLER, and IMP: an integrated modeling system. J Struct Biol 179, 269–278. 10.1016/j.jsb.2011.09.006

Zhang, Y., Liu, T., Meyer, C.A., Eeckhoute, J., Johnson, D.S., Bernstein, B.E., Nusbaum, C., Myers, R.M., Brown, M., Li, W., Liu, X.S., 2008. Model-based Analysis of ChIP-Seq (MACS). Genome Biol 9, R137. 10.1186/gb-2008-9-9-r137

Zhang, Z., Yu, W., Tang, D., Zhou, Y., Bi, M., Wang, H., Zheng, Y., Chen, M., Li, L., Xu, X., Zhang, W., Tao, H., Jin, V.X., Liu, Z., Chen, L., 2020. Epigenomics-based identification of oestrogen-regulated long noncoding RNAs in ER+ breast cancer. RNA Biology 17, 1590–1602. 10.1080/15476286.2020.1777769

