## Supplementary material for "Enhancer-driven local 3D chromatin domain folding modulates transcription in human mammary tumor cells": Image File-supl

### Fig1 sup- related to Fig 1

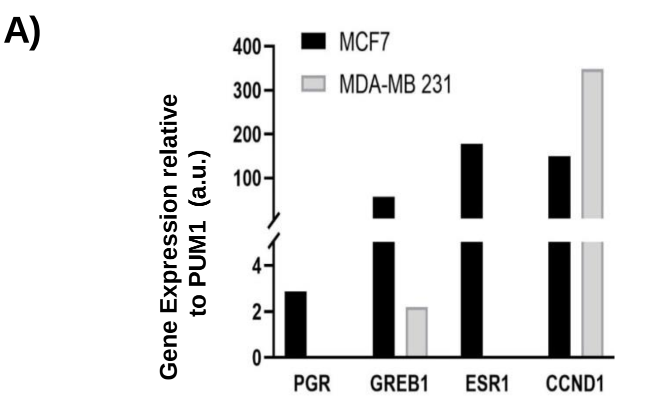


A) Expression levels of four ERa-regulated genes (PGR, GREB1, ESR1 and CCND1) confirmed by RT-qPCR in MCF7 and MDA-MB-231 cells. B) Interaction frequency 5C heat maps at 1.8 kb resolution surrounding the PGR gene showing two distinct TADs in MCF7 cells and erasing the TAD2 in MDA-MB-231 cells. C) Interaction frequency 5C heat maps at 1.8 kb resolution surrounding the ESR1 gene showing interactions between three ERBSs in MCF7 and their lost in MDA-MB-231 (green arrow), and more long-distance interactions in MDA-MB-231 cells (green arrowhead) compare to MCF7 profile. D) Interaction frequency 5C heat maps surrounding the control gene (PUM1) and the interaction counts (E) determined in MCF7 and MDA-MB-231 cells

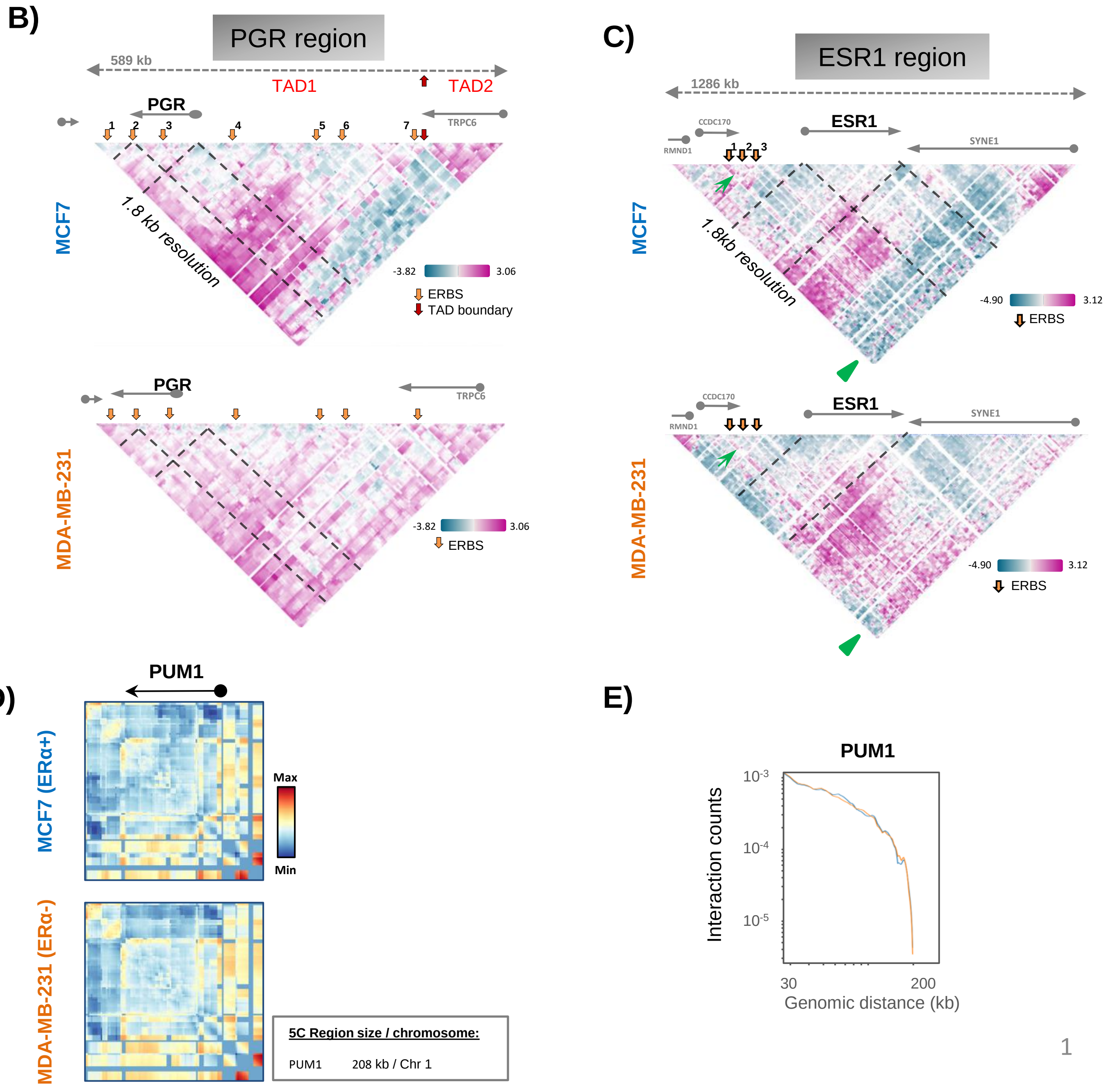

### Fig1 sup- related to Fig 1

F)

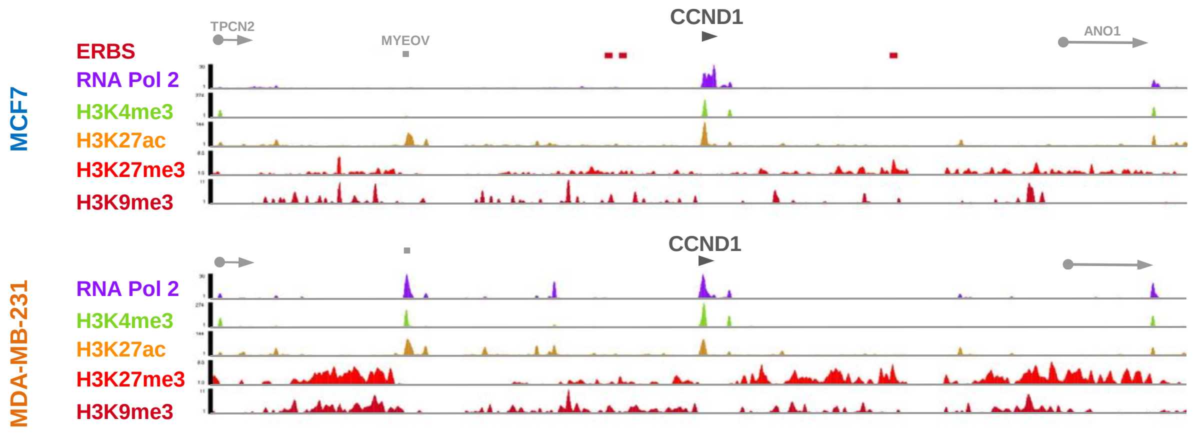


G)

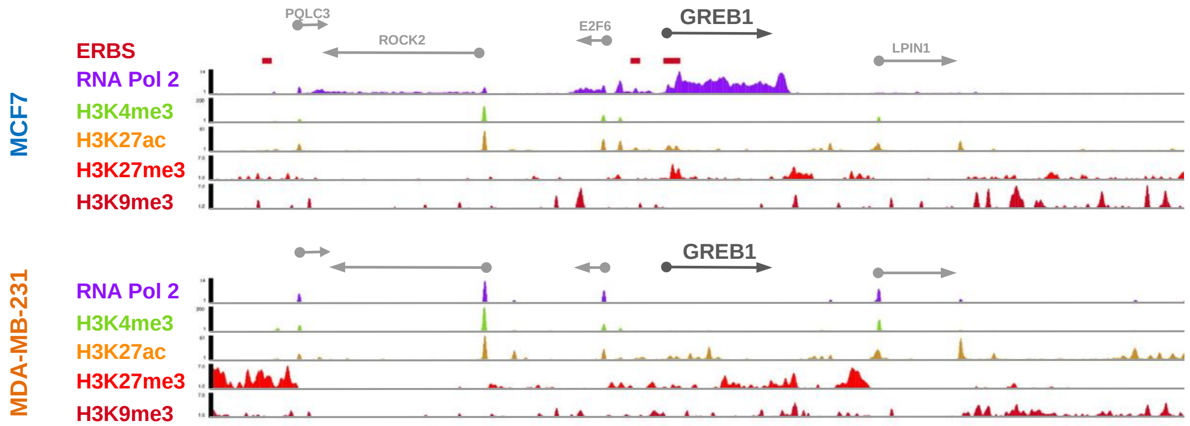


Chromatin landscape of the CCND1 (F) and the GREB1 (G) gene domains in MCF7 and MDA-MB-231 cells, from ENCODE (ENCODE Project Consortium, 2012; Guertin et al., 2014; Luo et al., 2020).

### Fig2 sup- related to Fig 3

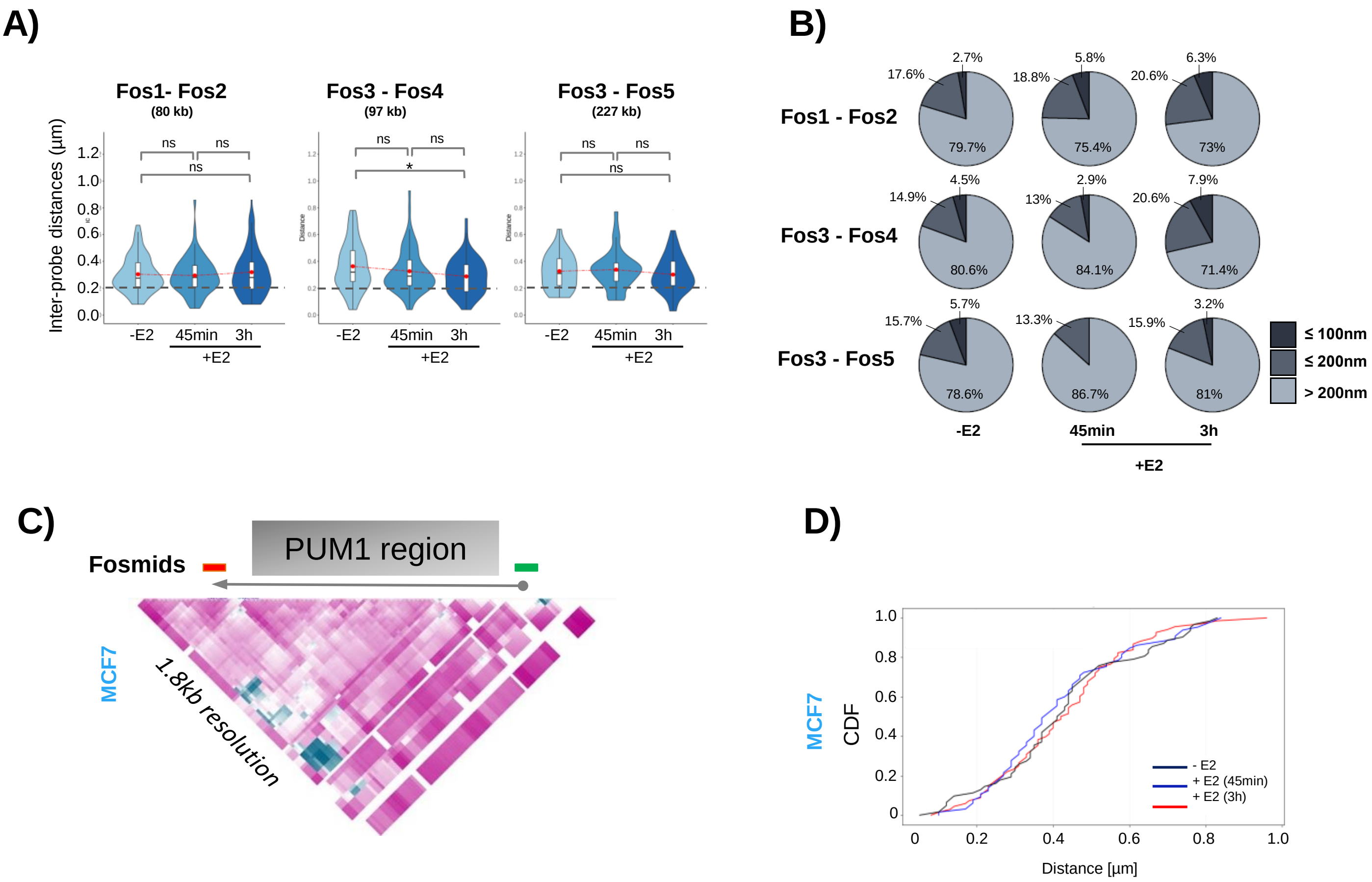


A) Violin plots representing inter-probe distances for different pairs of fosmids (n= 80-130 nuclei). Fisher's test: *P-values: >0.05 (ns), <0.05 (*), <0.01(**), <0.001 (***), <0.0001 (****). B) Pie charts illustrating the proportion of inter-probe distances > 200nm, interval from 200-100nm (≤ 200nm) and interval from 100-0nm (≤100 nm) for different pairs of fosmids in MCF7 cell stimulated or not by E2. Fisher's test was used for statistics. C) Genomic position of fosmid probes used for 3D DNA FISH analysis for control gene (PUM1). D) Cumulative distribution functions (CDF) for inter-probe distances obtained from 3D DNA FISH before (-E2) and after (+E2) transcription stimulation in MCF7 cells for PUM1.

### Fig 3 sup- related to Fig 4

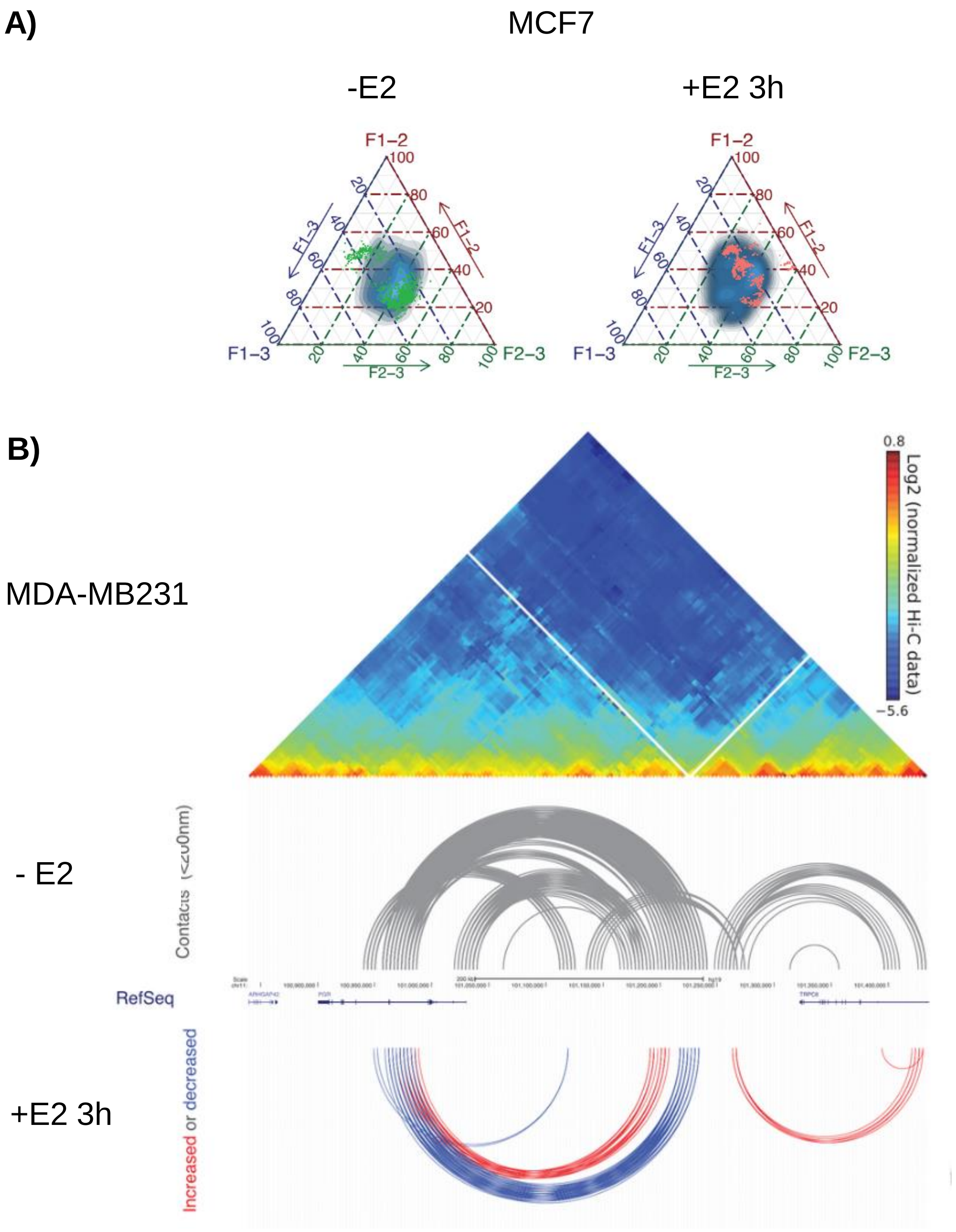


A) Comparison between three-way distances derived from 5C modeling (green and red dots, from A) and in situ 3D FISH measured distances (blue clouds, data from Fig. 3) in MCF7 before and after E2 treatment. The three axes show percentage of computed distances in all models or in all nuclei between genomic regions corresponding to selected fosmid sequences.

B) Contact frequencies in MDA-MB231 cells exposed to (+E2) or not (-E2) estradiol. Increased (red) or decreased (blue) in 50% of the cell population after 3h of estradiol exposure.

**Supplementary Table 1:** Datasets used in this study.

| Publication | Cell | Target | Data set | Time of E2 induction | E2 concentration |
|---|---|---|---|---|---|
| Li et al, 2019 (Li et al, 2019) | MDA-MB-231 | H3K4me3, H3K27ac, H3K27me3, H3K9me3, Pol II | GSE124379 | Untreated | - |
| ENCODE (Sloan et al., 2016) | MCF-7 | H3K4me3, H3K27ac, H3K27me3, H3K9me3 | H3K4me3 : ENCSR985MIB H3K27ac : ENCSR752UOD H3K27me3 : ENCSR761DLU H3K9me3 : ENCSR999WHE | Untreated | - |
| Kong et al. 2011 (Kong et al., 2011) | MCF-7 | GATA3/p300 | GSE29073 | 0 min, 45 min | 10 nM |
| Joseph et al. 2010 (Joseph et al., 2010) | MCF-7 | FOXA1, cFos, cJun | GSE26831 | 0 min, 3h | 10 nM |
| ENCODE (Sloan et al., 2016) | MCF-7 | MYC | MYC +E2 : ENCSR000DMP MYC -E2 : ENCSR000DMQ | 0 min, 45 min | 100 nM |
| Guertin et al. 2014 (Guertin et al., 2014) | MCF-7 | ER | GSE54855 | 0 min, 40 min, 2h40 | 100 nM |
| Zhou et al. 2019 (Zhou et al., 2019) | MCF-7 | H3K27ac | GSE108787 | 0 min, 1h, 4h | 100 nM |
| Maina et al. 2014 (wa Maina et al., 2014) | MCF-7 | PolII | GSE44800 | 0 min, 40 min, 2h40 | 10 nM |

Supplemental references:

Li, Kening, Congling Xu, Yuxin Du, Muhammad Junaid, Aman-Chandra Kaushik, and Dong-Qing Wei. 'Comprehensive Epigenetic Analyses Reveal Master Regulators Driving Lung Metastasis of Breast Cancer'. *Journal of Cellular and Molecular Medicine* 23, no. 8 (August 2019): 5415–31. https://doi.org/10.1111/jcmm.14424.

Guertin, M.J., Zhang, X., Coonrod, S.A., Hager, G.L., 2014. Transient Estrogen Receptor Binding and p300 Redistribution Support a Squelching Mechanism for Estradiol-Repressed Genes. Molecular Endocrinology 28, 1522–1533. https://doi.org/10.1210/me.2014-1130

Joseph, R., Orlov, Y.L., Huss, M., Sun, W., Li Kong, S., Ukil, L., Fu Pan, Y., Li, G., Lim, M., Thomsen, J.S., Ruan, Y., Clarke, N.D., Prabhakar, S., Cheung, E., Liu, E.T., 2010. Integrative model of genomic factors for determining binding site selection by estrogen receptor-α. Mol Syst Biol 6, 456. https://doi.org/10.1038/msb.2010.109

Kong, S.L., Li, G., Loh, S.L., Sung, W.-K., Liu, E.T., 2011. Cellular reprogramming by the conjoint action of ERα, FOXA1, and GATA3 to a ligand-inducible growth state. Mol Syst Biol 7, 526. https://doi.org/10.1038/msb.2011.59

Sloan, C.A., Chan, E.T., Davidson, J.M., Malladi, V.S., Strattan, J.S., Hitz, B.C., Gabdank, I., Narayanan, A.K., Ho, M., Lee, B.T., Rowe, L.D., Dreszer, T.R., Roe, G., Podduturi, N.R., Tanaka, F., Hong, E.L., Cherry, J.M., 2016. ENCODE data at the ENCODE portal. Nucleic Acids Res 44, D726-732. https://doi.org/10.1093/nar/gkv1160

wa Maina, C., Honkela, A., Matarese, F., Grote, K., Stunnenberg, H.G., Reid, G., Lawrence, N.D., Rattray, M., 2014. Inference of RNA polymerase II transcription dynamics from chromatin immunoprecipitation time course data. PLoS Comput Biol 10, e1003598. https://doi.org/10.1371/journal.pcbi.1003598

Zhou, Y., Gerrard, D.L., Wang, J., Li, T., Yang, Y., Fritz, A.J., Rajendran, M., Fu, X., Stein, G., Schiff, R., Lin, S., Frietze, S., Jin, V.X., 2019. Temporal dynamic reorganization of 3D chromatin architecture in hormone-induced breast cancer and endocrine resistance. Nat Commun 10, 1522. https://doi.org/10.1038/s41467-019-09320-9

**<u>Supplementary Table 2:</u>** Fosmids used in this study.

| Fosmid name | Whitehead (Sanger) name | Ensembl name | coordinates | | Size (bp) | midposition |
|---|---|---|---|---|---|---|
| Fos1 | *WI2-1069H1* | *G248P83181D1* | 100'857'188 | 100'893'758 | 36'570 (+) | 100'875'473 |
| Fos2 | *WI2-3198P11* | *G248P8001H6* | 100'930'139 | 100'972'364 | 42'226 (+) | 100'951'252 |
| Fos3 | *WI2-630J2* | *G248P80527E1* | 101'008'384 | 101'050'881 | 42'498 (+) | 101'029'663 |
| Fos4 | *WI2-2176P22* | *G248P87059H11* | 101'105'566 | 101'148'475 | 42'910 (+) | 101'127'021 |
| Fos5 | *WI2-758J11* | *G248P81257E6* | 101'238'076 | 101'275'616 | 37'541 (+) | 101'256'846 |
| PUM1 (3’) | *WI2-1091D18* | *G248P82995B9* | 31’359’975 | 31’403’626 | 43,652 (-) | 31’381’800 |
| PUM1 (5’) | *WI2-1757M9* | *G248P86664G5* | 31’540’133 | 31’581’406 | 41,274 (+) | 31’581’406 |
